# Reconsider phase reconstruction in signals with dynamic periodicity from the modern signal processing perspective

**DOI:** 10.1101/2020.09.29.310417

**Authors:** Aymen Alian, Yu-Lun Lo, Kirk Shelley, Hau-Tieng Wu

## Abstract

Phase is the most fundamental physical quantity when we study an oscillatory time series. There are many tools aiming to estimate phase, most of them are developed based on the analytic function model. Unfortunately, this approach might not be suitable for modern signals with *intrinsic nonstartionary structure*, including multiple oscillatory components, each with time-varying frequency, amplitude, and non-sinusoidal oscillation, e.g., biomedical signals. Specifically, due to the lack of consensus of model and algorithm, phases estimated from signals simultaneously recorded from different sensors for the same physiological system from the same subject might be different. This fact might challenge reproducibility, communication, and scientific interpretation and thus we need a standardized approach with theoretical support over a unified model. In this paper, after summarizing existing models for phase and discussing the main challenge caused by the above-mentioned intrinsic nonstartionary structure, we introduce the *adaptive non-harmonic model (ANHM)*, provide a definition of phase called *fundamental phase*, which is a vector-valued function describing the dynamics of all oscillatory components in the signal, and suggest a time-varying bandpass filter (tvBPF) scheme based on time-frequency analysis tools to estimate the fundamental phase. The proposed approach is validated with a simulated database and a real-world database with experts’ labels, and it is applied to two real-world databases, each of which has biomedical signals recorded from different sensors, to show how to standardize the definition of phase in the real-world experimental environment. Specifically, we report that the phase describing a physiological system, if properly modeled and extracted, is immune to the selected sensor for that system, while other approaches might fail. In conclusion, the proposed approach resolves the above-mentioned scientific challenge. We expect its scientific impact on a broad range of applications.

## 1. Introduction

Phase is the most fundamental physical quantity when we study the dynamics of a system of interest. It has been studied over centuries and is a standard material covered in almost all scientific fields. In a very general sense, we may collect all possible statuses of the system and call the collection the *phase space*, which might be a high dimensional space. The flow-volume loop [11] or pressure-volume diagram [12] are common examples that physicians commonly use in clinics for decision making. In the flow-volume loop example, the parameters of phase are the pressure and volume, which jointly describe the respiratory dynamics. In this paper, we focus on discussing the phase function of an oscillatory (or periodic) time series (or signal). Roughly speaking, the phase of an oscillatory time series is a value that describes the signal’s status within the span of a single oscillatory period (Figure 1a). It is closely related to the notion of frequency and period. In biomedicine, the phase contains crucial physiological information. For example, the pulse transit time reflects the phase relationship between electrocardiogram (ECG) and photoplethysmogram (PPG), and it is known to be related to the blood pressure [1, 2]; the coupling between the heart and lung, known as the cardiopulmonary coupling (CPC), is quantified by evaluating the phase relationship of the heart rate and respiration [3-5]; the coupling between different frequency bands of the electroencephalogram (EEG) or the amplitude-phase coupling [6, 7], has been well explored in the neuroscience society [7, 8], among others [9, 10]. This list of scientific merits of phase information is far from exhaustive.

**Figure 1:**
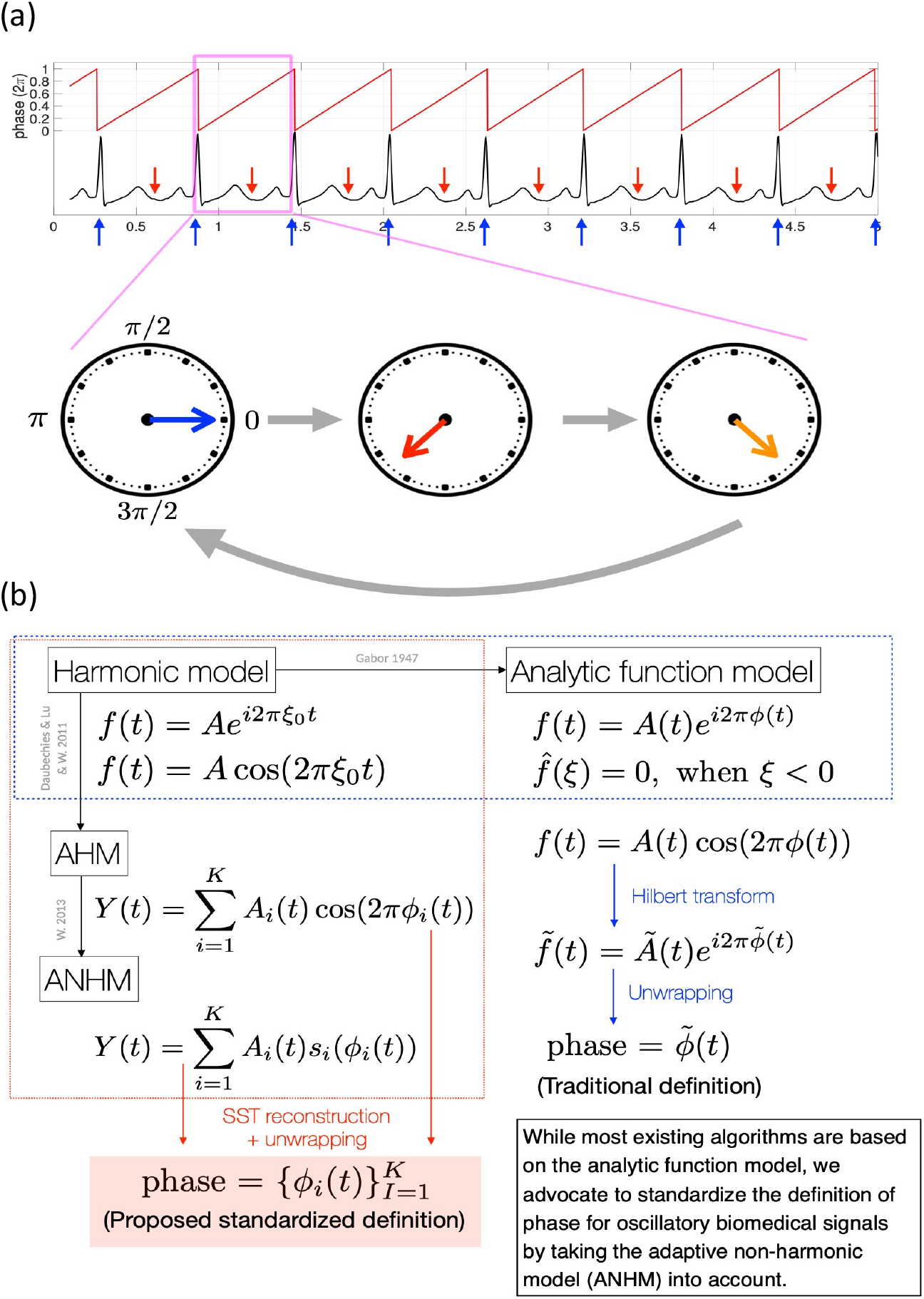
(a) The red curve on the top is the phase function associated with the ECG shown in the black curve, which ranges from 0 to 1 with the unit 2π. The cycle indicated by the magenta box is zoomed in to illustrate the idea of phase with a clock-like circling plot, where the time flow is indicated by the gray arrows. The blue arrows below the ECG signal indicate the status associated with the phase 0, and the red arrows above the ECG signal indicate the status associated with the phase 5π/_1_. The phase 0 is associated with the R peaks, and the phase 5π/_1_ is associated with the T end. In our proposed standardization, the phase is defined as a monotonically increasing function. This might cause some confusion in the illustrated phase in the red curve. Since the phase is the argument (or the angle) of the complex value, a phase value *θ* is not different from a phase value *θ* + 2*k*π for any integer *k*. Therefore, one way to plot the phase function is plotting the residue of the phase function by finding its value modulo 2π. (b) A summary of models for the definition of phase function. The traditional phase is shown on the right-hand side, and the proposed fundamental phase is shown on the left-hand side.

Historically, there has been rich literature discussing the theoretical foundation of phase function of a given time series. The approaches can be roughly classified into two categories based on how detailed we could model the signal. In the first category, when physiological knowledge is sufficient, it is possible to consider modeling the physical dynamical system by a set of differential equations and hence further describe and estimate the phase space and the phase. There is a huge literature discussing approaches in this category, ranging from the famous Hodgkin-Huxley model [13] to many current models like [14, 15], which again is a far from exhaustive list. While such a differential equation approach usually leads to an interpretable system, this direction is not the focus of this work since a satisfactory differential equation model for many real-world signals is usually lacking. For interested readers, we refer them to, for example, the monograph [16, 17] for more details.

In the second category, constructing a detailed model is challenging, and we count on signal processing techniques to define what a phase is. There has been a lot of work in this direction. From the model perspective, typical papers discussing the foundation of a phase function include, but not exclusively, [18-22]. The widely accepted model for the phase function is the *analytic function model* [18, 21, 23] proposed by Gabor back in 1946 [21], based on which most existing theories and phase extraction algorithms are developed. The Hilbert transform [24] with or without a bandpass filter design or its variations [25], time frequency (TF) analysis tools like continuous wavelet transform (CWT) [26] or others [27, 28] are commonly applied algorithms to estimate the phase function from an observed signal. This analytic function model and associated algorithms (hereafter called *traditional approach*) have been successfully applied to various scientific fields, like the communication system or information processing with solid theoretical support. In biomedicine, the traditional approach has usually been the choice [34], and it is claimed in Le Van Quyen M et al [35] that there are minor differences between the Hilbert transform and CWT on the intracranial or scalp EEGs. Recently, for the sake of handling nonstationary time series, algorithms like empirical mode decomposition (EMD) [31] and its variation like ensemble EMD (EEMD) [32] and variational mode decomposition (VMD) [33] have been introduced and widely applied. However, to our knowledge, despite empirical arguments, a theoretical support of these algorithms under a proper model is still missing, and we need to be careful when applying these tools (hereafter called *ad hoc approach* due to the lack of theoretical support). Based on the rich theoretical support from complex analysis, algorithms like Blaschke decomposition (BKD) [29] (or called adaptive Fourier transform [30]) based on the roots of an analytic function have also been considered recently. The potential of BKD has been shown before, but its development is still in its infancy.

While there have been many works in this direction, these approaches are limited. An ideal definition of phase function should not only be theoretically rigorous, but also be able to properly describe the status of the system with physical meaning. However, in biomedicine, depending on the signal, both the traditional and ad hoc approach based on the analytic model might not fulfill these criteria. For example, its application to the respiratory signal, PPG, peripheral venous pressure (PVP) [36], etc., might be problematic since they may contain multiple physiological oscillatory components, and each of them oscillates with *non-sinusoidal oscillatory pattern, time-varying frequency and amplitude*. The estimated phase might thus be not physiologically meaningful. See Figure 8(a) for an illustration (with more details in Section 5 below), where the phase estimated from the end-tidal CO_2_ (EtCO_2_) signal fluctuates fast so that the associated frequency goes beyond 0.8Hz, while the associated EtCO_2_ signal shown in Figure 2(a) obviously does not oscillate that fast. This leads to a problematic interpretation of what the phase is if it is determined by the traditional or ad hoc approach. Moreover, for the same system, different sensors lead to different oscillatory patterns. See Figure 2(b) for an illustration of four different respiratory signals recorded simultaneously. This diverse oscillatory pattern further impacts the estimated phase if we apply the traditional approach. See the discrepancy of the phases for the respiratory system estimated from different sensors by the traditional approach shown Figure 8(a) for an example. Clearly, such diverse information might lead to problematic scientific interpretation of what the status of the system is at each time.

**Figure 2:**
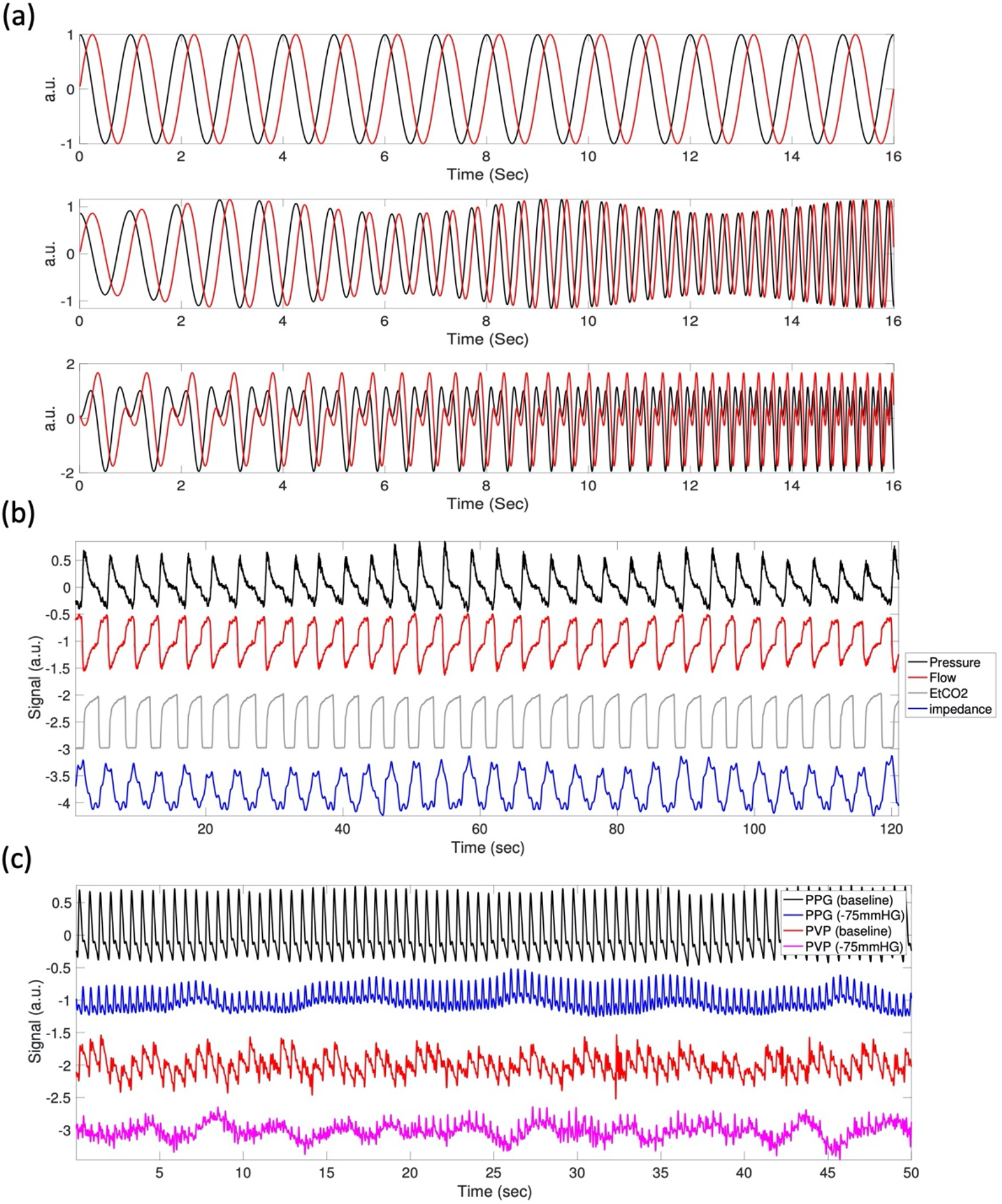
(a) illustration of simulated signals satisfying different models. Top: the harmonic model in equation (1) *f*(*t*) = *e*^*i*2 πt^; middle: the adaptive harmonic model in equation (3) 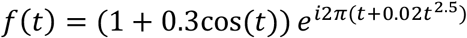; bottom: the wave-shape model in equation (5) with 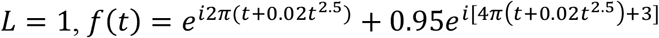. The black and red curves are the real and imaginary parts of the complex form, respectively. (b) An illustration of various respiratory signals recorded simultaneously. The morphologies of those respiratory signals are different, and this difference is mainly encoded in the multiples. (c) An illustration of photoplethymogram (PPG) and peripheral venous pressure (PVP) signals recorded simultaneously during the lower body negative pressure (LBNP) experiment. The four signals from top to bottom are the PPG signal during the baseline, the PPG signal during the -75 mmHg negative pressure, the PVP signal during the baseline, and the PVP signal during the -75 mmHg negative pressure. The morphologies of the same channels are different during different physiological status, and this difference is mainly encoded in the multiples.

In this paper, we focus on oscillatory time series with time-varying frequency and amplitudes and nontrivial oscillatory patterns, like respiratory signal and PPG, and argue that the traditional and ad hoc approaches might not be suitable for this kind of signals. This leads us to reconsider the notion of phase. We provide a solution by providing a new definition of phase called *fundamental phase* and propose proper tools for the analysis. To the best of our knowledge, this topic is less systematically discussed, except some reports [35]. The lack of consensus, or standard, might lead to the challenge of reproducibility, communication, and scientific interpretation.

The paper is organized in the following way. To address a broader audience, we start from layman’s language to review how an oscillatory biomedical signal is modeled and what is phase from the traditional perspective. This review leads to the suggested adaptive non-harmonic model (ANHM), which offers a platform to standardize the definition of phase that can be properly interpreted physiologically. The standardized phase is called the *fundamental phase*, since it is related to the fundamental component of each oscillatory component. We then suggest to estimate the fundamental phase by the time-varying bandpass filter (tvBPF) scheme based on time-frequency analysis tools, like the synchrosqueezing transform (SST) [37, 38], CWT [26] or BKD that will be detailed below (hereafter called the *proposed approach*). We illustrate how the algorithm works and compare it with different algorithms in a simulated dataset and a real-world database. In the end, we apply the proposed approach to resolve the phase standardization problem when we sense one system by different sensors simultaneously by applying the proposed approach to two real-world databases and comparing them with the traditional and ad hoc approaches. We shall mention that while we demonstrate the whole work with biomedical signals as examples, the model and algorithm can be applied to study signals outside biomedical society.

## 2. What is phase? From old to new

In this section, we discuss the notion of phase for oscillatory time series with different models. The non-oscillatory time series, particularly signals that are better modeled by a random process like electroencephalogram, is out of the scope of this paper, and we will discuss it in the Discussion section. See Figure 1(b) for an illustration of the relationship among these models and definitions.

### 2.1 Pure harmonic model

In the textbook setup [24], an oscillatory time series is usually modeled by the *pure harmonic model* that satisfies

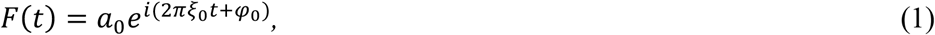

where *a*_0_ > 0 is the amplitude, *ξ*_0_ > 0 is its frequency, and *φ*_0_ ∈ [0,2π] is the *global phase shift*. Note that (1) is a typical example of the analytic model. The *phase function* is defined as 2π*ξ*_0_*t* + *φ*_0_. See Figure 2(a) for an example. Given F(t), it is transparent to evaluate the phase function directly by *unwrapping* the function 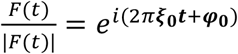. Usually, the signal we record is real-valued; that is, the signal is usually of the real part of F(t), and a common practice is applying the Hilbert transform to recover *F*(*t*) from its real part, and then get the phase function.

### 2.2 Adaptive harmonic model

However, most physiological signals are not harmonic since they oscillate with time-varying frequency and amplitude. For example, a subject might breathe at different rates at different moments [39]. This behavior can be modeled by:

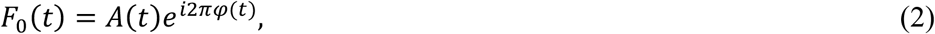

where *A*(*t*) is a smooth positive function and *φ*(*t*) is a smooth monotonically increasing function. We call *φ*^′^(*t*) > 0 the *instantaneous frequency* (IF) and *A*(*t*) the *amplitude modulation* (AM) of the function *F*_0_(*t*) [38]. In (2), the phase function we are concerned with in this paper is defined as *φ*(*t*). Note that (2) in general is not analytic and cannot be modeled by the analytic model. See Figure 2(a) for an example. Clearly, (1) is a special case of (3) when both the AM and IF are positive constants, and the phase is a linear function. Physically, at time *t*, the signal *F*_0_(*t*) repeats itself in about 1/*φ*′(*t*) seconds with an oscillatory magnitude *A*(*t*). This function has been shown to well approximate several physiological signals [36, 40]. In this model, since |*F*_0_ (*t*)| = *A*(*t*), the phase can be easily uniquely evaluated by unwrapping 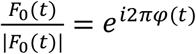.

### 2.3 Relationship between real and complex models and Vakman’s problem

While (2) in the complex formulation is commonly considered in the mathematical literature for the purpose of analysis, in practice, the signal is real. Thus, we consider the following model in the real formulation that is in parallel to (2),

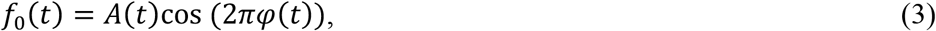

where the meanings of *φ*′(*t*), *φ*(*t*), and *A*(*t*) are the same as those in (2). It might be intuitive to define the phase function to be *φ*(*t*). However, this intuition needs a careful justification due to the *identifiability issue*. Indeed, it is easy to find infinite pairs of 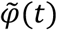, and *Ã*(*t*) so that 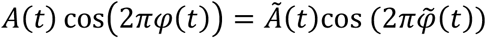 for all time *t*. Thus, without knowing *A*(*t*), the phase cannot be uniquely defined.

An intuitive approach to handle this identifiability issue is considering the Hilbert transform to convert (3) into an analytic function so that the phase can be uniquely defined. However, unlike the relationship between the pure harmonic model (1) and its real part, we face a challenge in this approach, which is closely related to the *Vakman’s problem* [41-43]. Specifically, applying the Hilbert transform to recover *F*_0_(*t*) from f_0_(t) is not usually mathematically possible since *F*_0_(*t*) is usually *not* an analytic function, nor a unitary function [18]. As a result, a spectral leakage is inevitable, which could be quantified in the global *L*^2^ sense [22, 44]. Therefore, the Hilbert transform cannot recover *F*_0_(*t*) unless some specific conditions are satisfied [22, 44]. However, these conditions are limited [18] and hold only for specific signals, like those considered in communication systems. We shall elaborate that even if the model is not mathematically correct, it could be *physically accurate* [18] and hence practically useful. A typical example is the frequency modulation with high carrier frequency in the radar communication. However, for the biomedical time series, this model might also be physically inaccurate since the carrier frequency is usually low compared with the variation of the varying frequency, and hence in general the phase defined under the model (2) cannot be estimated accurately or even correctly interpreted by the Hilbert transform. See Figures 4(a) and 6(a) for examples, and more discussions in the later section.

Despite the above limitations, it has been found that under some reasonable conditions, we can still make sense of the model (3) for the phase definition. A reasonable condition is the *slowly varying AM and IF* [37, 38]; that is, the AM and IF change slowly compared with the IF. Mathematically, this assumption is described by

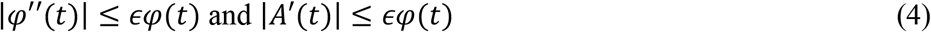

for some small *ϵ* > 0 for all time *t*. When (4) is satisfied, (3) is called the *intrinsic mode-type* (IMT) function. Under such assumption, it is shown in [37] that the identifiability issue is resolved, and the phase can be uniquely defined up to a negligible error. Thus, it does make sense to define the phase to be the same as that in the model (2). Note that (4) reflects physiological homeostasis – the variability from cycle to cycle does not change dramatically or randomly during a short time without external stimulations, while the system might migrate from one status to another over a long period.

### 2.4 More realistic model – multiple components with wave-shape function

There are more detailed features that cannot be captured by the IMT function. One feature is the *non-sinusoidal* oscillatory pattern. For example, ECG or PPG do not oscillate like a sine wave. See Figure 2(b,c) for example. Moreover, usually a biomedical signal is composed of several components with different physiological interpretations. For example, the PPG signal is composed of one hemodynamic component and one respiratory component known as the respiratory induced intensity variation (RIIV) [45]; the respiratory signal is composed of one respiratory component, and one hemodynamic component known as the cardiogenic artifact [46]. This fact holds for other biomedical signals. These observations lead to the ANHM [47, 48]:

#### Definition 1

*[Adaptive non-harmonic model (ANHM)]. A function f*(*t*) *satisfies the ANHM if it has the expression*

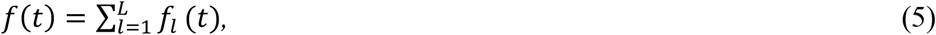

*where L* ≥ 1, *and for each l* = 1, …, *L*,

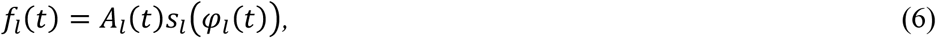

*where A*_*l*_(*t*) *and φ*_*l*_(*t*) *have the same meanings as those in (3) and satisfy (4), and s*_*l*_ *is a real 1-periodic function with mean 0, unit norm and* 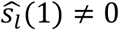. *We call f*_*l*_(*t*) *the l-th non-harmonic IMT function of the signal f*(*t*), *and s*_*l*_*(t) the wave-shape function (WSF) of the l-th oscillatory component*.

For example, for the respiratory signal, if the cardiogenic artifact exists [46], we have *L* = 2 in (5), where *f*_1_ models the respiratory signal and *f*_2_ models the cardiogenic artifact. See Figure 2(a) for a simulated example satisfying (5) with a non-sinusoidal oscillatory signal when *L* = 1, and Figures 2(b) and 2(c) for several real physiological signals with *L* = 1 in (5).

Clearly, (5) is a generalization of (3) since when *L* = 1 and *s*_*l*_(*t*) = cos (2π*t*), (5) is reduced to (3). This ANHM will be our base to propose a standardized definition of phase for a biomedical signal. Since our focus is the phase, we simplify the discussion without adding noise in (5), and refer readers to [37, 49] for the noisy cases. In practice, *L* in (5) might be known based on the background knowledge. When *L* is not known a priori, an estimation scheme is needed. While there exist some ad hoc approaches to estimate *L*, a systematic approach with theoretical support is so far lacking to our knowledge. Below, we assume the knowledge of *L*, and more discussion about *L* can be found in the Discussion section. We shall mention that there have been several generalizations, e.g. the time-varying WSF [48] and wave-shape manifold [50], to better capture the biomedical signals (see more in [36]). To simplify the discussion, we focus on Definition 1. We discuss how to define the phase function based on the model in Definition 1 in two cases: when *L* = 1 and when *L* > 1.

#### 2.4.1 *When L* = 1

In this case, the function satisfies *f*(*t*) = *f*_1_(*t*) = *A*(*t*)*s*(*φ*(*t*)), and *s*(*φ*(*t*)) is nothing but “stretching” *s*(*t*) by *φ*(*t*), and then “scaling” by *A*(*t*). We propose the following phase definition:

##### Definition 2

[*Fundamental phase when L* = 1]. *The phase function of f*(*t*) *satisfying (5) with L* = 1 *is φ*(*t*).

We need to mention a critical possible confusion point here, particularly when compared with the traditional approach to estimate the phase. Note that the Hilbert transform of *f*(*t*) can be written as 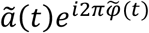. Thus, f(t) can be written as the real part of 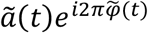, where if we enforced 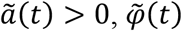 might *not* be a monotonic increasing function. In the traditional approach, the widely accepted definition of phase function of f(t) is 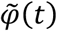, and hence the declared IF might be negative, which might be challenging to interpret. See a simulation in Figure 3 as an example of such a model and approach. Even worse, this Hilbert transform approach is sensitive to the input signal, even if two signals shown in Figure 3 are “visually similar”. From the complex analysis perspective, this fact comes from the “winding” effect caused by the non-sinusoidal oscillation.

**Figure 3:**
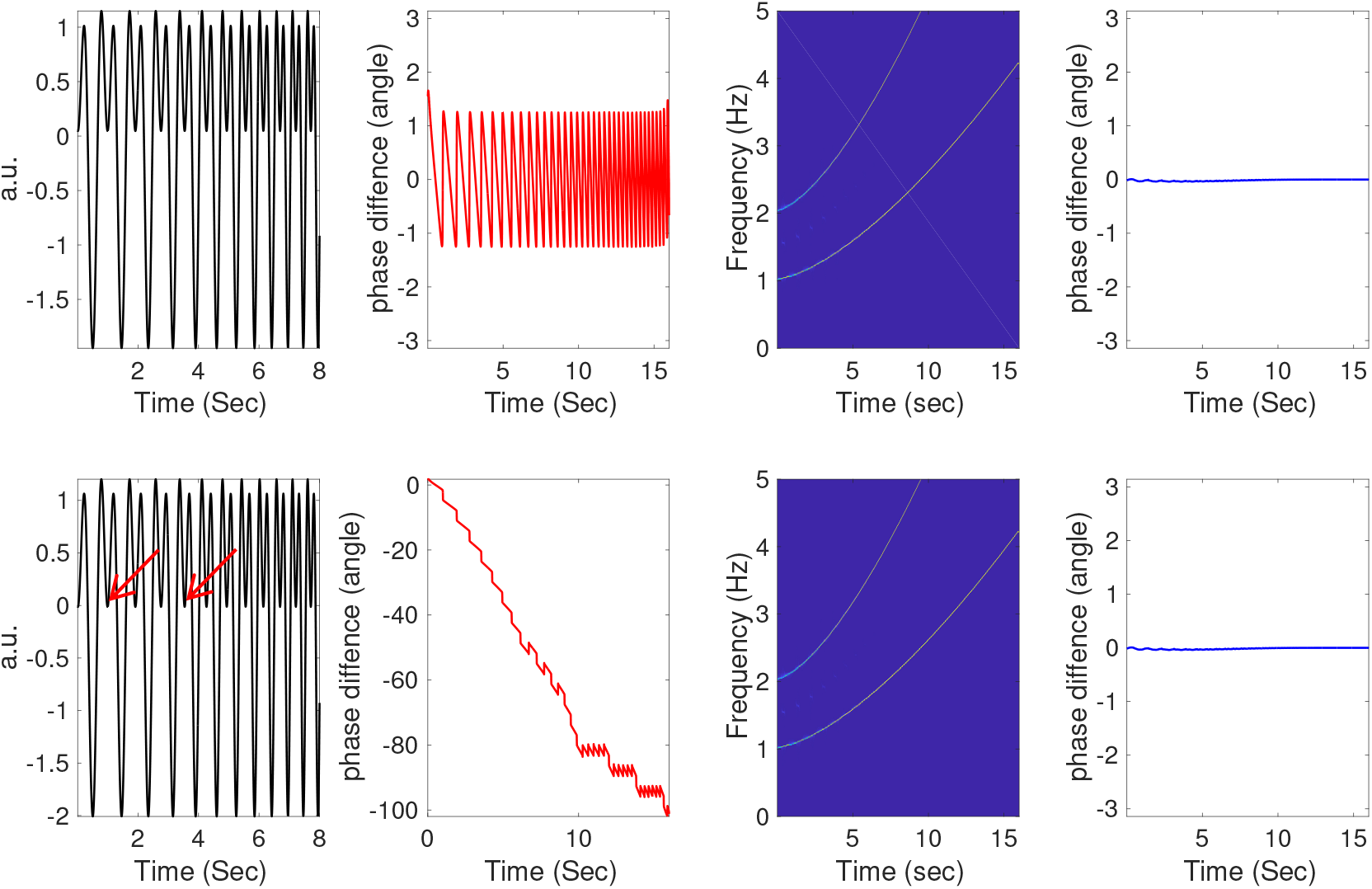
An illustration of the potential problem the Hilbert transform might encounter. The first signal, the real part of 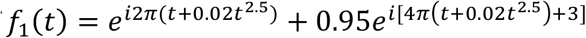, is shown on the left top panel, and the second signal, the real part of 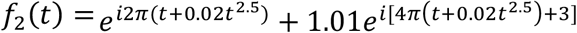 is shown on the left bottom panel. Note that the smaller trough of the real part of *f*_2_(*t*) crosses zero since the first harmonic has a larger amplitude than the fundamental component (red arrows). In the left middle columns, the red lines are the difference of the phase of the fundamental component, *φ*(*t*) = 2*π*(*t* + 0.02*t*^2.5^), and the estimated phase 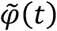 by directly applying the Hilbert transform. It is clear that for *f*_1_(*t*), there is an obvious zig-zag difference between the estimated phase function 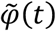 and the proposed phase function *φ*(*t*). However, for *f*_2_(*t*), there is an obvious cumulative phase estimation error as time evolves. This comes from the winding number issue due to the larger amplitude of the first harmonic. Due to the frequency modulation, the bandpass filter or continuous wavelet transform does not work as well (results not shown). In the right middle columns, the time-frequency representations of *f*_1_(*t*) and *f*_2_(*t*) determined by the synchrosqueezing transform (SST) are shown. In the right column, the blue lines are the difference of the estimated phase by SST and the proposed phase function *φ*(*t*). Despite the winding issue, SST provides a consistent phase estimation.

**Figure 4:**
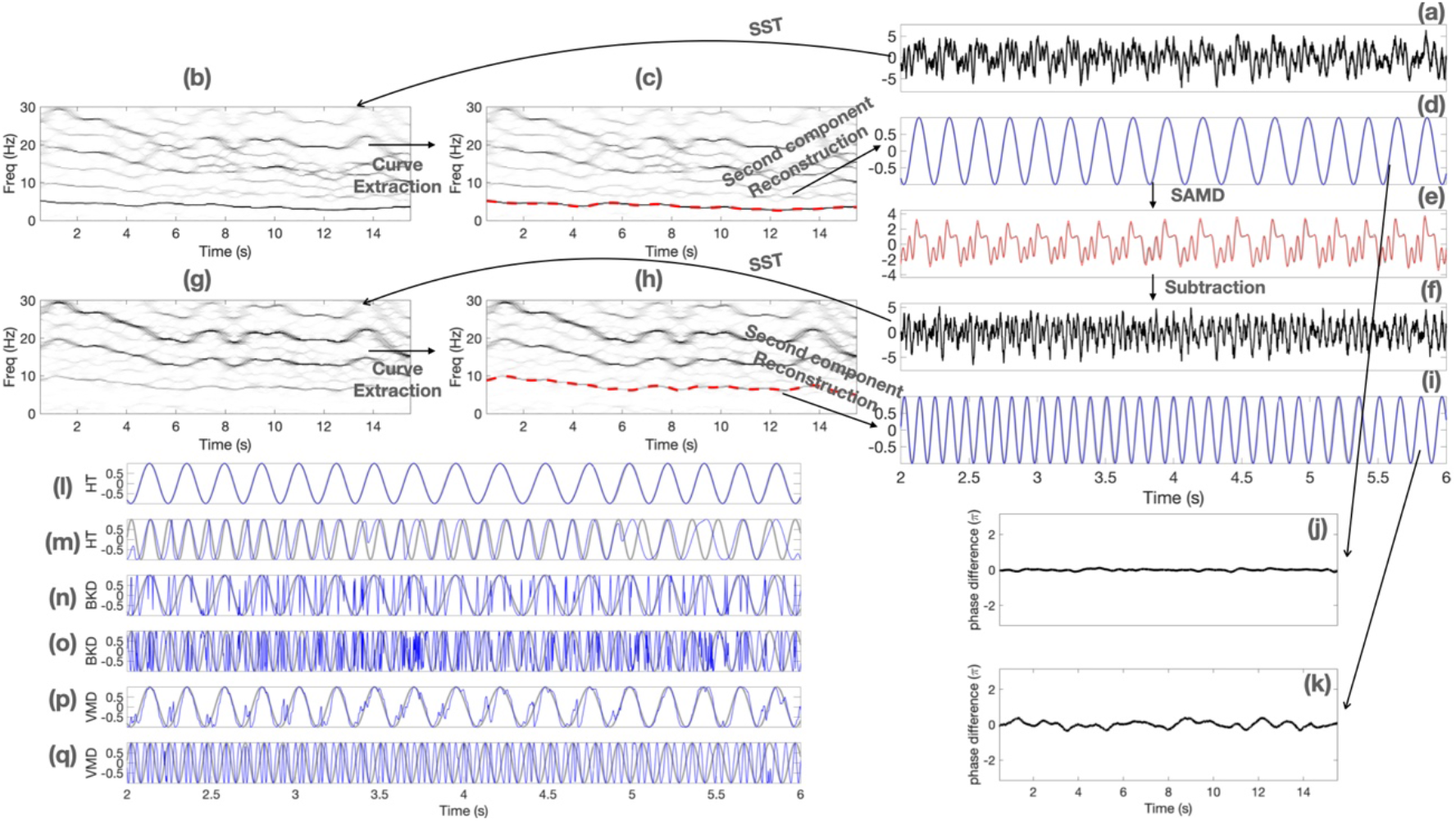
An illustration of the fundamental phase estimation by the proposed tvBPF with SST in a simulated database with ground truth, and the results of other methods, including the Hilbert transform with fixed band-pass filter (HT), the Blaschke decomposition (BKD) and the variational mode decomposition (VMD). (a) one realization of the clean signal *f* = *f*_1_ + *f*_2_ contaminated by noise, denoted as *Y*; (b) the time-frequency representation (TFR) of *Y*; (c) the extracted instantaneous frequency (IF) of *f*_1_ is superimposed as the red dashed curve; (d) the reconstructed fundamental component of *f*_1_ (blue) superimposed on the true fundamental component of *f*_1_ (gray); (e) the reconstructed *f*_1_, denoted as 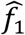 by apply the shape adaptive mode decomposition (red) superimposed on the true *f*_1_ (gray); (f) 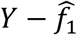, which is an estimate of *f*_2_; (g) the TFR of 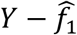; (h) the extracted IF of *f*_2_ is superimposed as the red dashed curve; (i) the reconstructed fundamental component of *f*_2_; (j) the difference of the estimated fundamental phase of *f*_1_ and the ground true; (k) the difference of the estimated fundamental phase of *f*_2_ and the ground true; (l)(n)(p): the estimated fundamental component of *f*_1_ by VDM, BKD and HT (blue) superimposed on the true fundamental component of *f*_1_ (gray); (m)(o)(q): the estimated fundamental component of *f*_2_ by VDM, BKD and HT (blue) superimposed on the true fundamental component of *f*_2_ (gray).

To look deeper into ANHM, consider a different interpretation of the signal *f*(*t*) = *f*_1_(*t*) = *A*(*t*)*s*(*φ*(*t*)). Note that the real signal in (5) could be expanded by the Fourier series of *s*, either pointwisely or in the distribution sense depending on the regularity of *s*, as

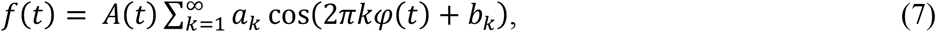

where we assume *a*_1_ > 0, and set *a*_*k*_ ≥ 0 for *k* > 1, *b*_*k*_ ∈ (0,2π], and 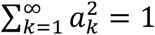. The interpretation of *f*(*t*) in (7) is different – *f*(*t*) is composed of multiple *sinusoidal* oscillatory components. We call *a*_1_*cos*(2π*φ*(*t*) + *b*_1_) the *fundamental component* and *a*_*k*_*cos*(2π*kφ*(*t*) + *b*_*k*_), *k* > 1, the (*k* − 1)*-th multiple* of the fundamental component. Note that the fundamental phase we consider, *φ*(*t*), is the phase function of the fundamental component, and this is the reason we call it the fundamental phase. One the other hand, the phase determined by the Hilbert transform approach contains information not only from the fundamental component, but also from *all* multiples. We shall also emphasize that sometimes researchers apply a bandpass filter (BPF) before applying Hilbert transform or CWT over a spectral band (i.e., traditional approach) to eliminate the impact of multiples. However, due to the time-varying frequency and amplitude, this spectral information of multiples might leak and contaminate the result. In short, while the model (5) could model various physiological time series, the traditional approach might not work as expected.

Now, we argue why it is better to define the phase to be *φ*(*t*) instead of using the Hilbert transform by taking physiological knowledge into account. Take ECG as an example (see Figure 1(a)), where the IF, AM and the WSF have their own physiological meanings. The WSF of ECG reflects the electrical pathway inside the heart, where the sensor is put, the heart anatomy, etc. Several clinical diseases are diagnosed by reading the WSF but not reading its Fourier series in (7). Moreover, while time moves forward, the physiological status should “move forward”; that is, the phase function should be monotonically increasing. This physiological fact suggests that it is better to separate the WSF from the phase function as in (6), which is precisely our definition of the phase function when *L* = 1. See Section 4 for concrete examples, like that in Figure 6(a).

Our definition has more benefits from the perspective of IF and AM. In the ECG example, it is well known that while the rate of the pacemaker is constant, the heart rate generally is not constant. The discrepancy comes from neural and neuro-chemical influences on the pathway from the pacemaker to the ventricle. This non-constant heartbeat rate could be modeled as the IF of the ECG. Moreover, the AM of the ECG reflects the breathing activity via the thoracic impedance. If we take model (7) into account, the interpretation of IF and AM might not be physiological.

#### 2.4.2 When L > 1

In this case, before providing the definition of “phase”, we shall discuss two facts. First, note that we may again consider to define the phase function via the Hilbert transform, which we also denote 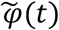. When *L* > 1, we may again have an interpretation problem even if *s*_*l*_(*t*) = cos (2π*t*) for *l* = 1, …, *L* since the phase information of different oscillatory components might be mixed-up. This leads to the same problem shown in Figure 3 discussed above. Take *f*(*t*) = cos(*φ*_1_(*t*)) + cos(*φ*_2_(*t*)) as a special example of (5) with *A*_1_(*t*) = *A*_*L*_(*t*) = 1 and *s*_1_(*t*) = *s*_2_(*t*) = cos (2π*t*), where we assume 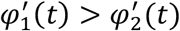. In this example, the phase 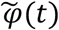 defined by the Hilbert transform will not be 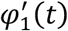 nor 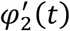 but a nonlinear interaction of 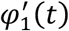 nor 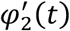.

Second, when *L* > 1, we claim that the phase of each non-harmonic IMT function shall have its own meaning, particularly when they represent different systems of interest. Take PPG as an example. The respiratory and cardiac components of PPG obviously have different physiological contents – one for the respiratory system dynamics and one for the hemodynamic system dynamics. So the phase 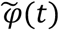 determined by Hilbert transform is a nonlinear mixed up of two systems, and it might not fully reflect the complicated physiology captured by PPG. These two facts lead us to define the phase of (5) to be a *collection* of phase function associated with each non-harmonic IMT component:

##### Definition 3

[*Fundamental phase when L* > 1]. *The phase of f*(*t*) *satisfying (5) when L* > 1 *is a set of phase functions* {*φ*_1_(*t*), …, *φ*_*L*_(*t*)}, *or equivalently a vector valued function Φ*(*t*) = [*φ*_1_(*t*), …, *φ*_*L*_(*t*)]^*T*^ *when t* ∈ ℝ.

Note that Definition 2 is a special case of the fundamental phase in Definition 3 when *L* = 1. Take PPG as an example again. Its fundamental phase, as a set of functions (or a vector-valued function) is a 2-dim vector valued function, where each dimension captures the dynamical information of one physiological system. In other word, by viewing the 2-dim vector valued function as two functions, each function describes the physiological dynamics of the associated physiological system. Compared with the traditional Hilbert transform approach, which always provides one function under the analytic model assumption, the fundamental phase respects the information hidden in each non-harmonic IMT component.

In summary, we propose to use ANHM in Definition 1 to describe various oscillatory physiological signals, and the associated phase function can be standardized and defined unambiguously with physiological meanings in Definitions 2 and 3, depending on the number of nonharmonic IMT functions. Moreover, we advocate that a phase function *φ*(*t*) of each non-harmonic IMT function shall capture the associated physiological status at each time, and the physiological status is described by the WSF.

## 3. Methodology

As we have discussed, traditional approaches might lead to problematic phase estimates when the oscillatory signal has a complicated structure, and these approaches may not be suitable to estimate the fundamental phase if the signal satisfies the ANHM. On the other hand, while it is possible to apply ad hoc approaches to estimate the fundamental phase, these ad hoc approaches have limited theoretical support and usually they do not perform well (see the simulation and real-world databases below for details). Motivated by the need to estimate the fundamental phase from the signal with theoretical supports, we need different methods. In this section, we mention two theoretically solid modern methods that could be applied to estimate the fundamental phase, including

1. the *time-varying bandpass filter (tvBPF)* with CWT [26] and SST [38];
2. BKD.

CWT and SST have been well developed in the time-frequency analysis society with solid theoretical support. To apply them to extract the fundamental phase, the key step is tvBPF that respects the non-constant IF. Below, we will quickly summarize CWT and SST and focus on how to carry out tvBPF to estimate the fundamental phase given in Definition 3. BKD, on the other hand, is a new technique that is based on complex analysis. Compared with CWT and SST that deeply depend on the Fourier theory, BKD is based on the root distribution of the signal inside the unit disk on the complex domain. So BKD can be understood as a nonlinear Fourier transform technique that respects the nonconstant IF.

The Matlab code and dataset used in Section 4 can be found in [51] and the Matlab implementation of BKD can be found in [58].

We shall emphasize that any tools that can separate and reconstruct the fundamental component of each non-harmonic IMT function could be used to estimate the fundamental phase, while in this paper we only discuss CWT, SST and BKD to avoid distraction. The reason we consider CWT is that it is widely applied in the biomedical signal processing field. We consider SST since it is a novel algorithm suitable for this purpose and has been widely applied recently. BKD is a novel signal processing tool that is based on a different mathematics theory and has potential to be applied to handle the phase problem.

### 3.1 tvBPF with CWT and SST

In this section, we first summarize CWT and SST, and then describe the tvBPF scheme to estimate the fundamental phase.

#### 3.1.1 Summary of CWT and SST

Fix the Gaussian function 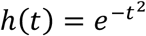. While the window can be more general, we focus on the Gaussian function to simplify the discussion. Take the signal *f*, and assume it is a tempered distribution. The short time Fourier transform (STFT) [27, 28] of *f* is defined as

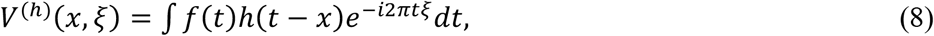

where *x* ∈ ℝ means time and *ξ* > 0 means frequency. We call the set of pairs(*x, ξ*) the time-frequency (TF) domain, and *V*^(*h*)^ is the TF representation (TFR) of *f*(*t*) via STFT. Call |*V*^(*h*)^(·,·)|^2^ the *spectrogram*.

CWT is different from STFT while sharing several similarities. Consider an analytic mother wavelet *ψ*, which has mean zero and unit norm, and we assume *ψ* has sufficient smoothness, integrability and admissibility [26] to simplify the discussion. The CWT of *f*(*t*) is defined as

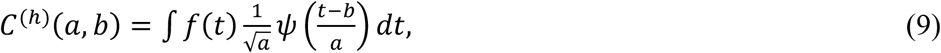

where *b* ∈ ℝ means time and *a* > 0 means the scale. Call |*C*^(*h*)^(·,·)|^2^ the *scalogram*. Note that *ψ* has mean 0 and unit norm, so it is oscillatory. But stretching *ψ* by *a*, we could “fit” the signal and detect how fast the signal oscillates. Note that *ψ* is oscillatory since it has mean 0. Thus, the scale *a* captures the inverse of frequency up to a proper multiplicative constant. Clearly, CWT and STFT are different. In STFT the signal is truncated by a fixed window and the truncated signal is analyzed by the Fourier transform, while in CWT, a pattern (mother wavelet) is stretched to fit the signal. See [27, 28] for a detailed treatment of the difference.

STFT and CWT can be classified as the linear-type TF analysis tools. SST, on the other hand, is a nonlinear-type TF analysis defined on STFT [52] or CWT [26], where the nonlinearity comes from a nonlinear transform of the TFR determined by STFT or CWT. To demonstrate the idea, we focus on SST defined on STFT, and the idea of SST defined on CWT is the same. The main observation of SST is that for each (*x, ξ*), *V*^(*h*)^(*x, ξ*) is a complex value, and its *complex angle* contains useful information that leads to SST. Define the *reassignment rule* as

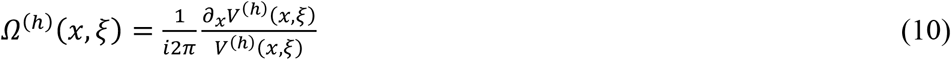

when *V*^(*h*)^(*x, ξ*) is nonzero and −∞ otherwise. With the reassignment rule, the SST of *f* is defined by reallocating STFT coefficients by

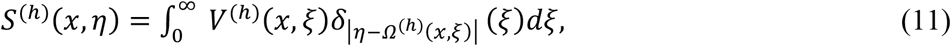

where *η* > 0 is the frequency and *δ* denotes the Dirac measure supported at 0. Compared with the STFT, the main benefit of SST is enhancing the contrast of the TFR determined by STFT by sharpening the spectrogram and hence alleviating the possible spectral mix-up between different components. We may call |*S*^(*h*)^|^2^ the *time-varying power spectrum* since it provides momentary spectral information.

The key idea beyond SST is that the reassignment rule tells us how fast the signal oscillates at each time. Indeed, for a given pair of *x* and *ξ*, if we rewrite 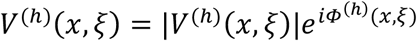, where *Φ*^(*h*)^(*x, ξ*) ∈ [0,2π], and assume that |*V*^(*h*)^(*x, ξ*)| > 0, we have *log* (*V*^(*h*)^(*x, ξ*)) = *log*(|*V*^(*h*)^(*x, ξ*)|) + *iΦ*^(*h*)^(*x, ξ*). Since 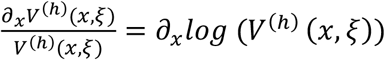 and *Φ*^(*h*)^(*x, ξ*) encodes “how fast” the signal oscillates at time *x*, when the magnitude *log*(|*V*^(*h*)^(*x, ξ*)|) does not vary too fast, the reassignment rule gives us the information about how fast the signal oscillates at time *x*. Thus, the squeezing step (11) sharpens the TFR. The main benefit of sharpening the TFR is enhancing the contrast, so that we can more accurately extract the IF and hence more accurately reconstruct the non-harmonic IMT components for the fundamental phase estimation [67, 68]. For readers with interest in theoretical results and statistical inference by SST, we refer them to [37, 38, 49].

#### 3.1.2 Numerical implementation of CWT and SST

Suppose the signal *f*(*t*) is sampled *N* times uniformly with the period Δ*t* > 0; that is, the *i*-th sample is *f*(*i*Δ*t*). The CWT implementation is standard, and we refer readers to, e.g., [53, 54]. On the other hand, SST is implemented by discretizing (8), (10) and (11) directly. In (8) and (10), the frequency axis is sampled from −1/(2Δ*t*) to 1/(2Δ*t*) with the resolution Δ*ξ* = 1/(*N*Δ*t*) following the Nyquist rate rule. In (11), the frequency axis η is sampled from 1/(*N*Δ*t*) to 1/(2Δ*t*) with the resolution 1/(*N*Δ*t*). But in practice, we may choose a coarser grid (a lower resolution) to speed up the computational time; that is, the frequency axis η is sampled from Δη to *m*Δη, where Δη > 0 is the resolution and *m* ∈ ℕ satisfying *m*Δη = 1/(2Δ*t*). Denote the discretized TFR *S*^(*h*)^ as a *N* × *m* matrix ***S***.

#### 3.1.3 *tvBPF when L* = 1

The tvBPF is composed of two steps, which are common for CWT and SST. To simplify the discussion, we explain the tvBPF with SST and assume that the fundamental component dominates to simplify the discussion (this condition can be removed by applying the de-shape algorithm [48], but since this topic is tangential to the notion of phase, we omit it here to avoid distracting the readers). We start from *L* = 1. First, we estimate the IF of the fundamental component by the ridge extraction [23]:

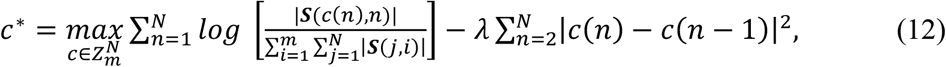

where *λ* > 0 is a parameter chosen by the user and *Z*_P_ = {1,2, …, *m*}. Clearly, *c* is a *N*-dim vector. The first term in (12) means we fit a curve *c* to the TFR ***S*** so that the curve captures the most intense pixels in the TFR, while the second term imposes the smoothness constraint on the curve *c*. By balancing between these two terms, *c*^*^ represents a smooth curve on the TFR that captures the IF of the fundamental component. In this work, we choose *λ*=0.5. According to the established theory [37, 38], the IF of the fundamental component at time *l*Δ*t* is *c*^*^(*l*)Δη. Second, we reconstruct the fundamental component of *f*(*t*) = *f*_1_(*t*) = *A*(*t*)*s*(*φ*(*t*)) from ***S*** via

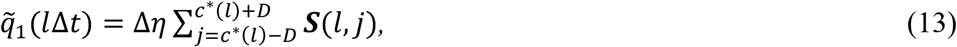

where *D* ∈ ℕ is chosen by the user. That is, at time *l*Δ*t*, we sum up the frequency band centered at the IF *c*^*^(*l*)Δη, with the bandwidth DΔη. See Figure 7(a) for an illustration, where the dashed curves below and above the solid red curve indicate the band, while the solid red curve indicates the extracted ridge. In practice, *D*Δη is suggested to be about 1/10 of the fundamental frequency. In this work, we choose *D* so that *D*Δη = 0.1.

The nomination of tvBPF comes from (13) since the frequency band may vary from time to time. It is thus like filtering the signal so that the filtered signal 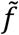 has different spectral bands at different times.

#### 3.1.4 *tvBPF when L* > 1

When *L* > 1 in (5), the fundamental phase is obtained by iterating the following steps. Apply the tvBPF in Section 3.1.2 to the signal *f*(*t*) and get the fundamental phase of the first non-harmonic IMT function, denoted as 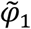, via setting 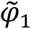 by unwrapping 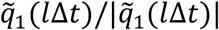. Similarly, we have an estimate of *A*_1_(*l*Δ*t*)*a*_1,1_ via evaluating 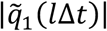. With the estimation of the fundamental phase of the first non-harmonic IMT function and its associated AM information, we apply the shape-adaptive mode decomposition (SAMD) [68] to reconstruct the first non-harmonic IMT function *f*_1_(*t*), denoted as 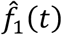. Then, subtract 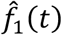 from *f*(*t*) and obtain a new function denoted as *f*^(1)^(*t*). Then, apply the tvBPF in Section 3.1.2 to *f*^(1)^(*t*) and obtain an estimation of the second non-harmonic IMT, denoted as 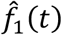, and its associated fundamental component 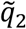. By iterating these steps *L* times, we obtain 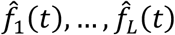 and the associated fundamental components 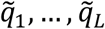.

#### 3.1.5 Obtain the fundamental phase

With the estimated fundamental components of all non-harmonic IMT components, denoted as 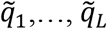, we estimate the fundamental phase defined in Definition 3. According to the established theory [52], for 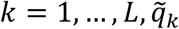 well approximates the complex form of the fundamental component of the *i*th non-harmonic IMT component; that is, for *f*_*k*_(*t*) = *A*_*k*_(*t*)*s*_*k*_(*φ*_*k*_(*t*)), we have 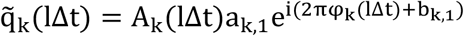 up to a negligible error. According to (6), since *A*_*k*_(*l*Δ*t*)*a*_*k*,1_ is not zero, the fundamental phase at time *l*Δ*t*, {*φ*_1_(*l*Δ*t*), …, *φ*_*L*_(*l*Δ*t*)} can be estimated by unwrapping 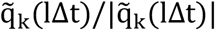 for *k* = 1, …, *L*. The accuracy of these estimators is guaranteed by the established theory [52].

#### 3.1.6 A comparison with the traditional approaches

The key idea of tvBPF is respecting the fact that the time-varying IF could lead to a spectral leakage, which is challenging to be captured by one fixed spectral band when multiple components exist or when the WSF is non-trivial. However, the traditional approach usually depends on the BPF with a *fixed* spectral band, even if CWT or other TF analysis tools are applied. In other words, the time-varying IF might lead to a broad spectral band, so that the spectral contents of different components or the harmonics of the non-trivial WSF overlap and hence a fixed spectral band fails to capture the desired information. This issue can be visualized in Figure 3. In that Figure, we illustrate the estimated phase of a simulated simple chirp signal with non-sinusoidal oscillation. Clearly, the phase estimated by the Hilbert transform is far away from the fundamental phase. Also, we see that due to the time-varying IF, the IF of the second harmonic in the beginning overlaps the IF of the fundamental component in the end. Therefore, a BPF with a fixed spectral band cannot isolate the fundamental component. These troubles we encountered by the Hilbert transform are eliminated if we apply the tvBPF scheme proposed above. As we will show below in the real-world data, even if the IFs of different components have separate ranges, it is not always a good idea to apply the traditional approach, when the signal oscillates with non-constant IF and AM.

### 3.2 Blaschke decomposition (BKD)

BKD is essentially different from CWT and SST mentioned above from the theoretical perspective, but it also captures the notion of tvBPF. To summarize BKD, a quick recap of the Fourier series in the complex analysis language is helpful. Take a holomorphic function *F*: ℂ → ℂ. We may trivially write it as *F* = *F*(0) + (*F*(*z*) − *F*(0)). Since 0 is a root of *F*(*z*) − *F*(0), we have *F*(*z*) − *F*(0) = *zF*_1_(*z*) for another holomorphic function *F*_1_(*z*). By iterating this procedure, that is, we unwind one zero at each iteration, we get

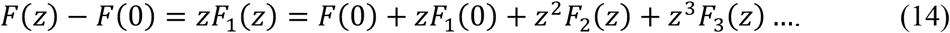

If we set *z* = *e*^*it*^, we have recovered the Fourier series of the function *F* restricted on *∂*𝔻, where 𝔻 is the unit disc on the complex domain. The key observation made by the authors in [29] is that we can generalize the idea of Fourier series by unwinding *all* zeros of *F* at once by factoring out a normalized product of all zeros. This decomposition is a classical object in complex analysis known as the *Blaschke factorization* [55], which states in the following way. Any holomorphic function *F*: 𝔻 → ℂ can be written as *F*(*z*) = *B*(*z*)*G*(*z*), where *G* has no roots inside the unit disk 𝔻 and *B* has magnitude 1 on *∂*𝔻 satisfying

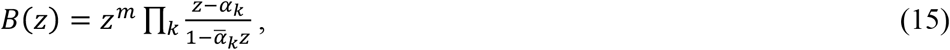

where *α*_*k*_ are roots of *F* inside the unit disk. Usually, *B* is called the Blaschke product. An application of this fact repeatedly generalizes Fourier series and allows us to decompose the function *F* as

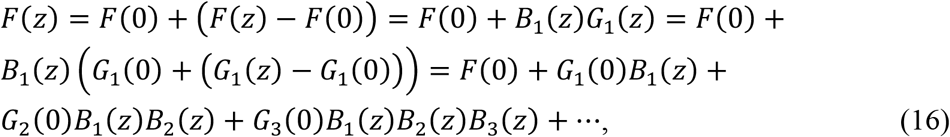

where the step *F*(*z*) − *F*(0) = *B*_1_(*z*)*G*_1_(*z*) is the Blaschke factorization collecting all roots inside the unit disk in *B*_1_(*z*), so that *G*_1_(*z*) is free of roots inside the unit disk. Similar explanation holds for the other decomposition steps. Denote

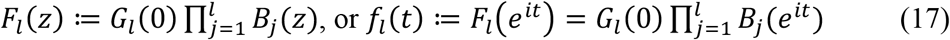

to be the *l*-th factored component by the above decomposition. Mathematically, *F*_*l*_(*z*) has a very natural physical interpretation. First, *G*_*l*_(0) is understood as the amplitude of the *l*-th component, and the complex angle of 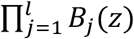 describes the frequency of the *l*-th component. Indeed, if we set the Blaschke product

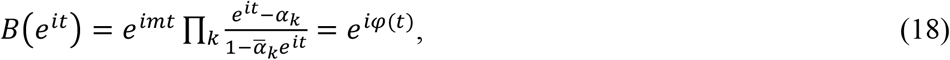

then by a direct calculation we have

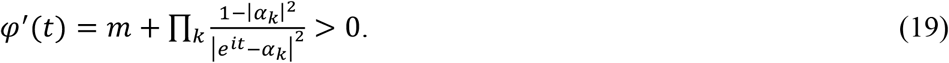

This peculiar property says that the complex angle of the Blaschke product, *φ*(*t*), is always monotonically increasing, so that physically *φ*^′^(*t*) is always positive and could be understood as the IF. This decomposition of *F*(*z*) into *F*_1_(*z*), *F*_2_(*z*), … is called the BKD, which is clearly a direct generalization of the Fourier series in the sense that we take *all* roots but not just 0. By construction, the IF of *F*_*k*_(*z*) is higher than *F*_*l*_(*z*) when *k* > *l*. Clearly, due to the nonlinear dynamics of the roots involved in the BKD algorithm, it is categorized as a nonlinear-type TF analysis. Compared with SST, BKD is developed based on the analytic model. It can be shown that the ANHM is closely related to the analytic model [56]. Thus, BKD is potential to be applied to extract the fundamental phase. We refer readers to [29, 56, 57] for more theoretical details.

#### 3.2.1 Numerical implementation of BKD

The numerical implementation of BKD is based on a well-known theoretical result. While theoretically we need to know the roots at each step, numerically this knowledge is not needed, thanks to the theoretical result reported in [4]. In fact, since |*F*| = |*G*| and *ln*|*G*| is analytic in the disk, by denoting 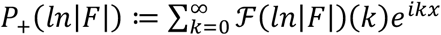, where ℱ indicates the Fourier series, we have formally that *G* = exp (*P*_+_(ln (|*F*| + *ε*))) and *B* = *F*/*G*. Therefore, *G* can be computed easily by applying the Fast Fourier Transform ℱ(*ln*|*F*|)(*k*) instead of finding all roots of *F*, where *ε* > 0 is a small constant to stabilize the blowup of natural log evaluation when |*F*| is close to zero. This BKD implementation is numerically stable and efficient, and a variety of numerical experiments have been carried out in [29] that validate these numerical properties.

#### 3.2.2 Obtain the fundamental phase by BKD

When *L* = 1 in the ANHM, the desired fundamental phase can be extracted by BKD by the phase of *F*_1_(*z*). To guarantee that the estimated phase accurately, the WSF should have a dominant fundamental component; otherwise, the phase will be contaminated by the phases of harmonics due to the winding effect. See [56, 57] for more discussions. Thus, when *L* = 1, BKD should be applied only when the oscillation is close (but can be different from) the sinusoidal oscillation.

When *L* > 1 and the WSF is sinusoidal, the fundamental phase can be estimated by the phases of *F*_1_(*z*), …, *F*_*L*_(*z*). To guarantee that the estimated phase is accurate, *F*_*l*_(*z*) should dominate the strength of the summation of *F*_*l*+1_(*z*), …, *F*_*L*_(*z*); otherwise, the phase will be contaminated by the phases of other components due to, again, the winding effect. Thus, when *L* > 1, BKD should be applied only when the oscillatory patterns are sinusoidal, and the strengths of IMT components decay sufficiently fast.

When *L* > 1 and the WSFs are general, based on our knowledge, it is so far not clear what is the best condition to guarantee if we can obtain the fundamental phase, and how.

While there are some known limitations, we consider BKD a potential tool to extract fundamental phase since it is theoretically guaranteed under some conditions. How to handle these theoretical problems is of interest on its own, but it is out of the scope of this paper and will be explored in our future work. While we will demonstrate results based on BKD, we must warn the users that

### 3.3 Illustration of the algorithm

We illustrate how to estimate the fundamental phase with simulated signals. To have a more real simulation, we consider the smoothened Brownian path realizations to model the AM and IF of the non-harmonic IMTs [66]. Denote *W* to be the standard Brownian motion defined on [0, ∞). Then, define the smoothened Brownian motion with bandwidth B > 0 as Φ_*B*_ ≔ *W* * *K*_*B*;_, where *K*_*B*_ is the Gaussian function with the standard deviation B > 0 and * denotes the convolution operator. Given T > 0 and parameters *ζ*_1_, …, *ζ*_6_> 0, we then define the following family of random processes on [0, *T*]:

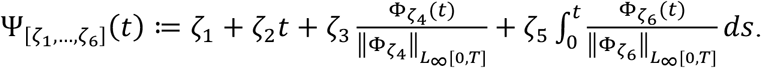

The simulated signal consists of two components, *f*_1_(*t*) and *f*_2_(*t*), (i.e. *L=2*) on [0,16], where *A*_1_(*t*), *A*_2_(*t*), *φ*_1_(*t*), and *φ*_2_(*t*) are independent realizations of Ψ_[2,0,1,2.5,0,0]_(*t*), Ψ_[1.,7,0,1,2,3,0,0]_(*t*), Ψ_[0, π,0,0,1,6,2,2]_(*t*) and Ψ_[0,8,0,0,3,2]_(*t*) respectively, and the associated WSFs, *s*_1_(*t*) and *s*_2_(*t*), are independent realized of 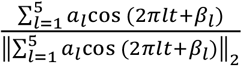, where *a*_1_ follows the 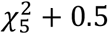 distribution, *la*_*l*_ follows the 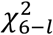 distribution for *l* = 2, …, 5, and *β*_*l*_ follows the uniform distribution over [0, 2π] and *a*_*l*_ and *β*_*l*_ are independent. We focus on those realizations that 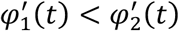 and 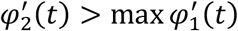; that is, when the spectral band of both non-harmonic IMT functions overlap. The signal is sampled at 512 Hz and a 10dB Gaussian white noise is added, where the noise level is defined as 10*log*_10_[VAR(signal)/VAR(noise)] with VAR indicating the variance. We also show the results with other methods. Since the phase of the signal determined by the Hilbert transform is a single function which is not comparable, we instead consider the fixed BPF approach with the Hilbert transform, where the spectral band for *φ*_1_(*t*) is determined by the range of 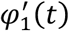 and that for *φ*_2_(*t*) is by the range of 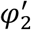(*t*); that is, the ground truth is used to avoid the technical issue of determining the spectral band. For a fair comparison, we use the same spectral band for the curve extraction in (12) when we apply the tvBPF approach. We also consider BKD mentioned above, and an ad hoc approach VMD. See Figure 4 for one realization of the simulated signal, and the steps of estimating the fundamental phase of the signal, *φ*_1_(*t*), and *φ*_2_(*t*), by different methods, where the proposed tvBPF approach we demonstrate is based on SST, and the results of different methods. We see that the due to the non-trivial WSFs, the signal is complicated and hard to see the oscillatory pattern. Even under this situation, the fundamental phase can be accurately estimated by the proposed tvBPF approach (Figure 4(j,k)), but not accurate with other methods. Note that the TFR determined by SST is very complicated (Figure 4(b)) due to the interaction of non-constant IFs, nontrivial WSFs and multiple components. After removing the estimated first non-harmonic IMT function, the TFR determined by SST is cleaner and associated with *f*_2_(*t*) (Figure 4(g)). The phase difference of the estimated fundamental phase and the ground truth is shown Figure 4(j,k). An ideal phase estimation should be close to 0 all the time. An interesting fact is that by the traditional fixed BPF approach, the estimation of *φ*_1_(*t*) is more accurate compared with that of *φ*_2_(*t*) (Figure 4(p,q)). Specifically, the estimated fundamental component associated with *f*_2_(*t*) is deformed. This come from the fact that the spectral band around *φ*_2_(*t*) is contaminated by the harmonics of *φ*_1_(*t*), while the spectral band around *φ*_1_(*t*) is not impacted by anything. The results by BKD and VMD are not ideal, while we can see that the “speed of oscillatory” is captured by both methods (Figure 4(l-o)).

Next, we quantify the accuracy of the estimation, and compare it with other approaches. We independently realize the signal and noise for 100 time, and evaluate the root mean square error (RMSE) of the estimated fundamental phases. The median ± median absolute deviation of 100 RMSEs of estimating *φ*_1_(*t*) by the proposed tvBPF based on SST, the Hilbert transform approach with fixed BPF, BKD and VMD are 0.25 ± 0.2_1_, 0.31 ± 0.27, 1.27 ± 0.26 and 1.29 ± 0.33 respectively, and the RMSE of estimating *φ*_2_(*t*) are 0.30 ± 0.27, 0.98 ± 0.32, 1.62 ± 0.08 and 1.68 ± 0.02 respectively. The performance of the proposed tvBPF approach is significantly better than others if tested with the Wilcox rank test with the significant level set to 0.05 with Bonferroni correction. Note that the proposed approach is robust to noise, and this robustness has supported by existing theoretical results [37, 49]. Due to the iterative process of the proposed tvBPF scheme, it is expected that the estimation of *φ*_2_(*t*) is worse than that of *φ*_1_(*t*).

Then, we show that BKD could work better when the WSFs are sinusoidal. Indeed, we consider the same smooth Brownian model like the above with *A*_2_(*t*) being the realization of Ψ_[0.85,0,0.5,2.3,0,0]_(*t*) and *a*_1_ = 1 and *a*_*l*_ = 0 for *l* = 2, …, 5 for both WSFs. Note that we intentionally force the amplitude of the second non-harmonic IMT function to be smaller to fulfill the currently known theory of BKD. The overall RMSE results with 100 realizations for estimating *φ*_1_(*t*) by the proposed tvBPF based on SST, the Hilbert transform approach with fixed BPF, BKD and VMD are 0.05 ± 0.11, 0.12 ± 0.05, 0.19 ± 0.08 and 0.30 ± 0.11 respectively, and the RMSE of estimating *φ*_2_(*t*) are 0.12 ± 0.11, 0.90 ± 0.18, 0.88 ± 0.16 and 1.70 ± 0.08 respectively. Again, the performance of the proposed tvBPF approach is significantly better than others if tested with the Wilcox rank test with the significant level set to 0.05 with Bonferroni correction. Note that in this case, BKD performs much better compared with the case when the WSFs are non-sinusoidal with statistical significance. Moreover, for 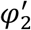, its median is better than that of the Hilbert transform with fixed BPF but without statistical significance. On the other hand, the Hilbert transform approach with fixed BPF performs very well for 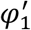, but less ideal for 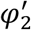, which again comes from the spectral overlap. In summary, we know that the non-trivial WSF plays a significant role when we estimate the fundamental phase.

Next, we demonstrate how the algorithm is applied to a real PPG signal and explain what information the fundamental phase can offer. We consider the CapnoBase IEEE TBME Respiratory Rate Benchmark database^1^ that consists of 42 recordings of 8 mins long from static patients under spontaneous or controlled breathing. The recordings include end-tidal CO_2_ (EtCO_2_), ECG and PPG signal, and all are sampled at 300 Hz. Labels from an expert are available for the timestamps of R peaks from ECG and inspiration initiation from EtCO_2_ as the ground truth. We use these experts’ labels to determine the ground truth phase in the following way. Suppose there are *L* timestamp labels in EtCO_2_ by experts at time *t*_*k*_, *k* = 1, …, *L*. The ground truth respiratory phase is defined by fitting the nonuniform samples 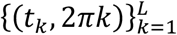. The same procedure is applied to the ECG signal to determine the ground truth cardiac phase. The resulting fundamental phase estimation and a comparison with other methods is shown in Figure 5. First, note that the curves associated with the respiratory component do not exist in Figure 5(c) since the respiratory component has been removed. We can clearly see that the fundamental phase, as a set of two functions, captures the cardiac and respiratory dynamics separately (Figure 5(f,g,h)). Specifically, in Figure 5(f-g), the reconstructed fundamental components of the first and second non-harmonic IMT functions coincides with the EtCO_2_ and ECG oscillations respectively. This fact is further confirmed by evaluating the difference between the estimated fundamental phase and the ground truth phase shown in Figure 5(h). Note that an ideal phase estimation should be close to a constant all the time, where the constant represents the global phase shift between the estimated fundamental phase and the ground truth phase. The results by Hilbert transform with a fixed bandwidth are also reasonable since the respiratory component and cardiac component have distinct spectral range and the harmonics of the respiratory component are not strong, while the results by BKD and VMD are not ideal due to the non-trivial WSFs. To quantify the performance of these algorithms, we carry out the analysis over all 42 cases. Since one case has a broken EtCO_2_ signal, it is removed from the analysis. The overall RMSE results over 41 cases for estimating the fundamental phase of the respiratory component by the proposed tvBPF based on SST, the Hilbert transform approach with fixed BPF, BKD and VMD are 0.37 ± 0.35, 0.63 ± 0.33, 1.73 ± 0.12 and 1.77 ± 0.18 respectively, and the RMSE of estimating the cardiac component are 0.15 ± 0.14, 0.22 ± 0.15, 1.70 ± 0.12 and 1.65 ± 0.28 respectively. The performance of the proposed tvBPF approach is significantly better than others if tested with the Wilcox rank test with the significant level set to 0.05. In summary, the proposed fundamental phase of the PPG signal captures the dynamic information of the cardiac and respiratory systems, which is potential for the further analysis, like the variability analysis. Moreover, the proposed tvBPF approach could estimate the fundamental phase from the PPG signal accurately.

**Figure 5:**
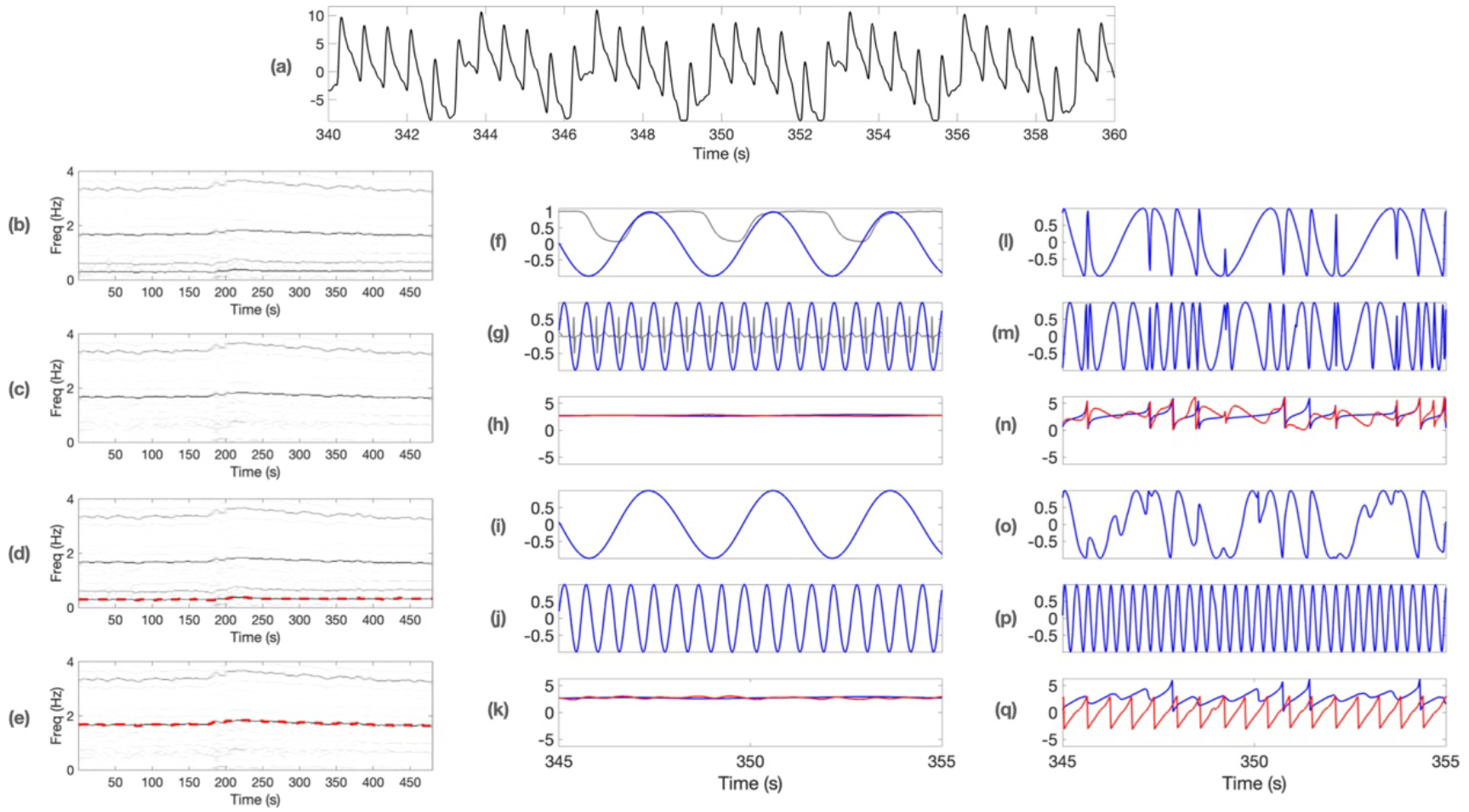
An illustration of the fundamental phase estimation by the proposed tvBPF with SST in a photoplethysmogram (PPG) signal (shown in (a)) with experts’ labels on the simultaneously recorded electrocardiogram (ECG) and end-tidal CO_2_ (EtCO_2_) signals. The experts’ labels lead to the ground truth phases of the respiratory and cardiac dynamics. The results of other methods, including the Hilbert transform with fixed band-pass filter (HT), the Blaschke decomposition (BKD) and the variational mode decomposition (VMD) are also shown. (b,d) the time-frequency representation (TFR) of the PPG signal shown in (a), where the extracted instantaneous frequency (IF) of the respiratory component is superimposed in (d) as a red curve. (c,e) the time-frequency representation (TFR) of the PPG signal subtracted by the reconstructed respiratory component, where the extracted instantaneous frequency (IF) of the cardiac component is superimposed in (e) as a red curve. (f,g) the reconstructed fundamental components of the respiratory and cardiac components are shown in blue curves, with the simultaneously recorded ECG and EtCO_2_ superimposed as gray curves respectively. (h) the phase differences between the estimated fundamental phases and the ground truth phases for the respiratory and cardiac components are shown in blue and red curves respectively. (i-k) the same as (f-h) but generated by HT. (l-n) the same as (f-h) but generated by BKD. (o-q) the same as (f-h) but generated by VMD.

**Figure 6:**
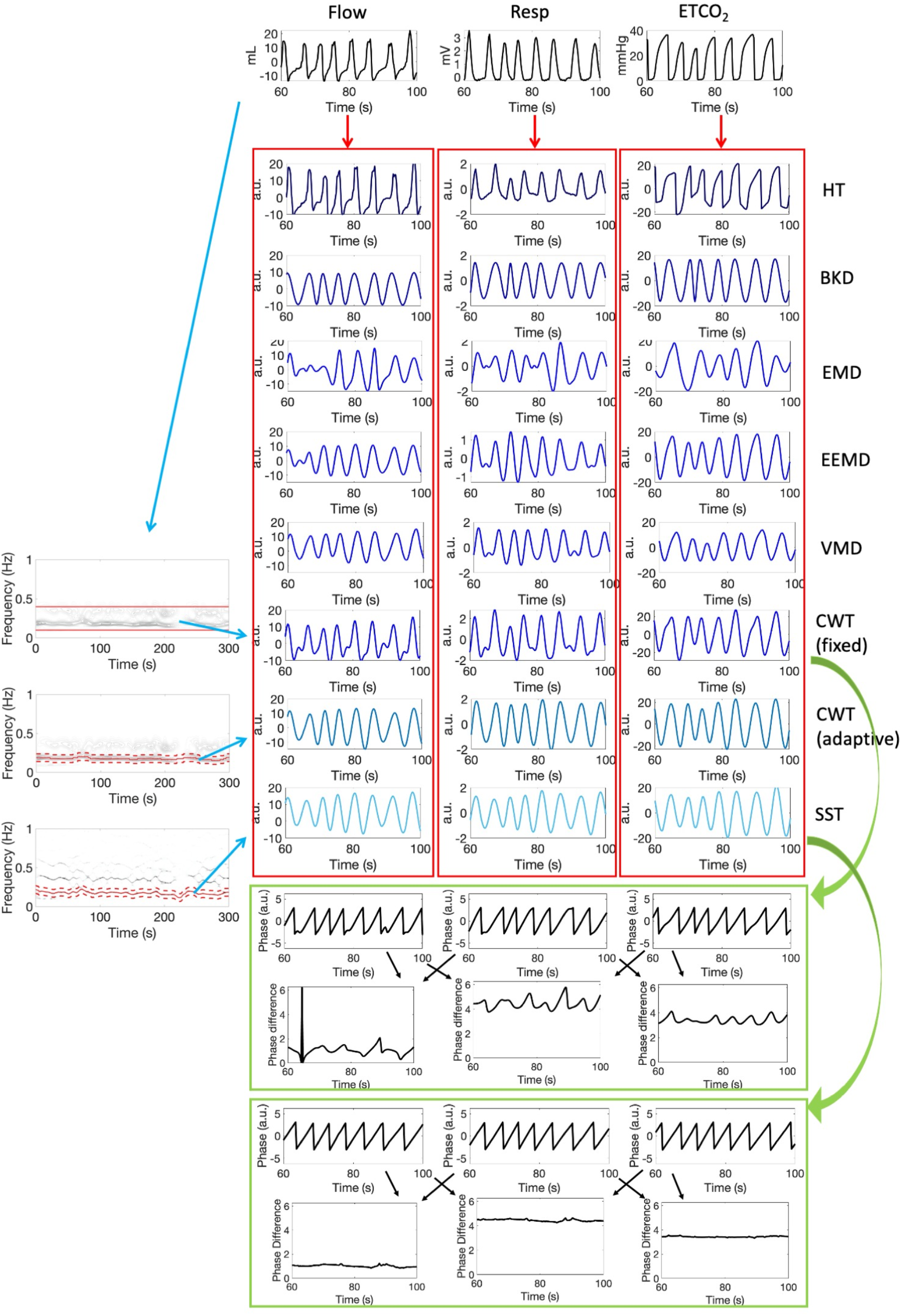
A comparison of different phase reconstruction methods from different respiratory sensors. (a) Three different respiratory signals. (b) The time-frequency representation (TFR) of the flow, impedance and end-tidal CO_2_ (EtCO_2_) signals, which were determined by CWT (top and middle) and SST (bottom) respectively. The fixed spectral band for CWT (CWT-fixed) reconstruction is shown in the top subfigure as red lines. The time-varying spectral bands for CWT and SST are plotted as dashed red curves above and below the solid red curve indicating the instantaneous frequency in the middle and bottom plots. (c) The reconstructed fundamental component by different methods. Each column is associated with one signal, and each row is associated with one method. (d) The phase estimated by CWT-fixed mod by π is shown in the top row, where the phases are different. The phase difference between two phases estimated from two sensors is shown in the bottom row. (e) Same as (d), but the phase is estimated by SST. It is obvious that the fluctuation is less compared with the CWT-fixed method. The phase estimated by CWT with the time-varying bandpass filter is similar (not shown).

**Figure 7:**
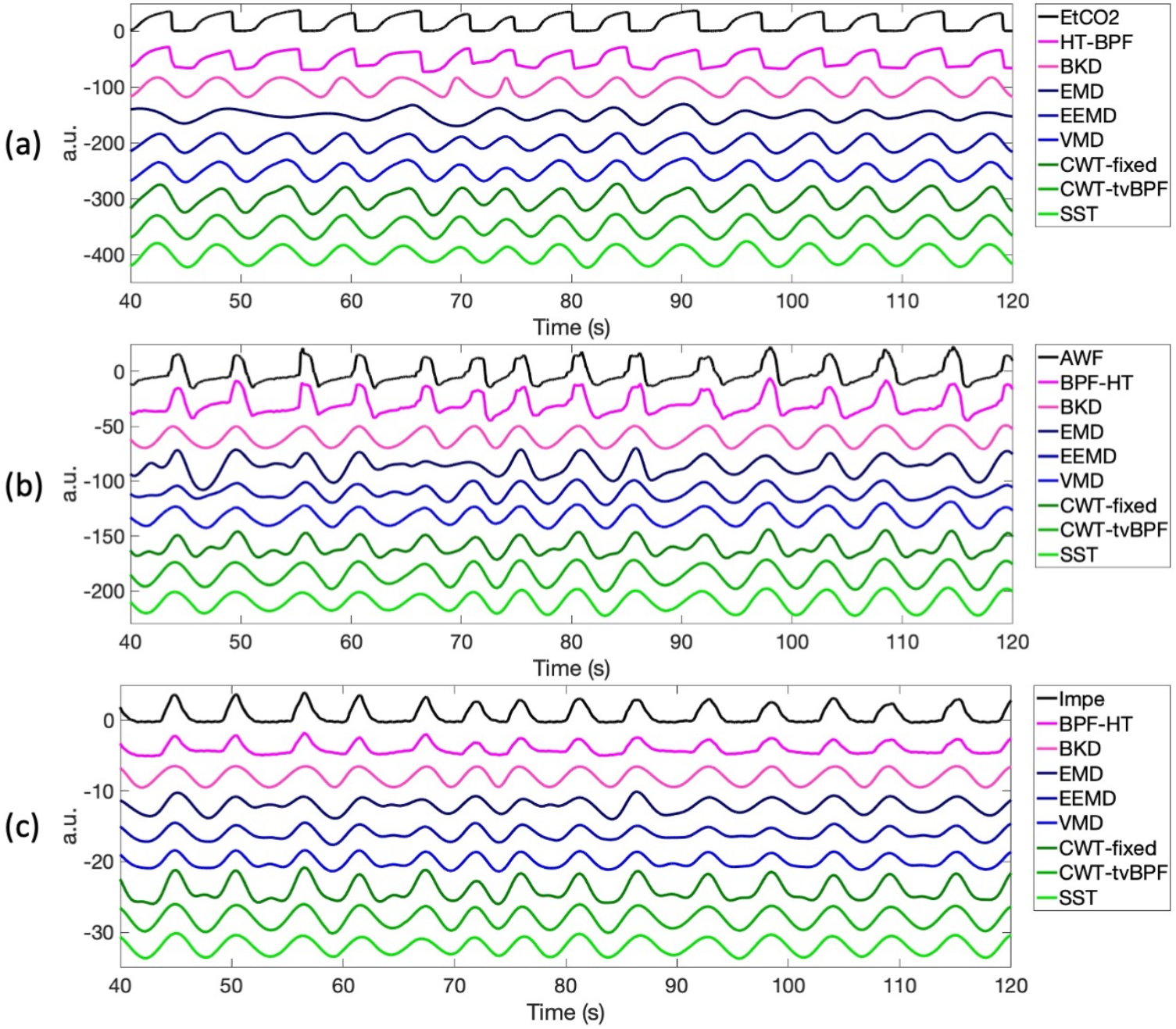
Demonstration of decomposed components for the standard phase estimation by different algorithms. (a) is the end-tidal CO2 (EtCO2) signal, (b) is the flow signal (AWF), and (c) is the impedance (Impe) signal.

**Figure 8:**
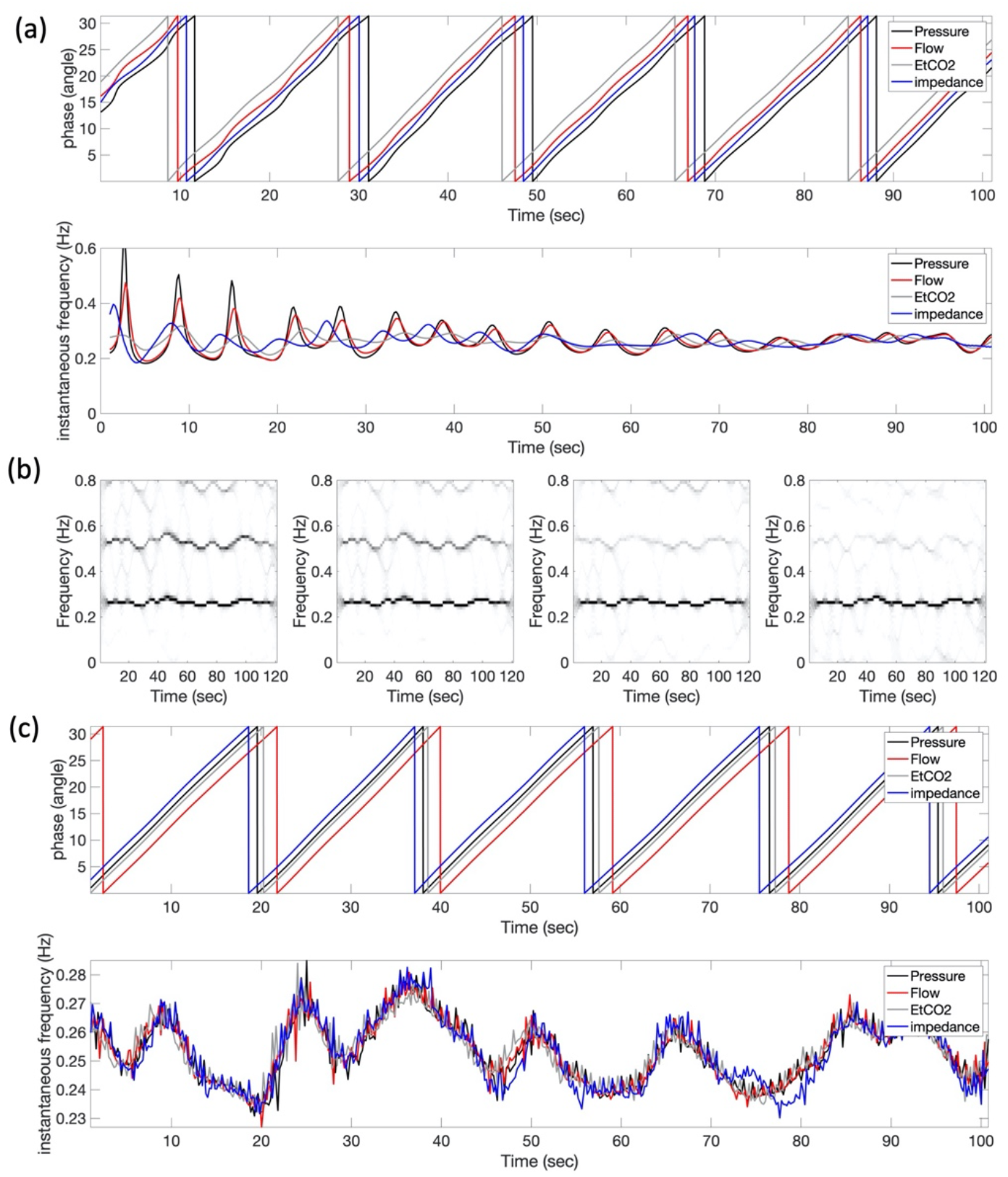
Demonstration of two typical algorithms. (a) An example of determining phases of various respiratory signals by the Hilbert transform followed by the band-pass filter (BPF-HT), where the band is 0.1-0.4 Hz determined by the dominant respiratory frequency. The phases (plotted with a mod of 10π to enhance the visualization), and hence their derivatives, are not consistent from one sensor to another. More importantly, while the definition of instantaneous frequency is mathematically correct, physiologically it is not sensible. The result by the continuous wavelet transform with a fixed spectral band (CWT-fixed) is similar and not shown. (b) An example of determining phases of various respiratory signals by the synchrosqueezing transform (SST). From left to right: the time-frequency representations of the pressure, flow, endtidal CO_2_, and impedance signals determined by SST. (c) Top panel: the phases of four respiratory signals estimated by SST (plotted with a mod of 10π to enhance the visualization). Bottom panel: the instantaneous frequency of four respiratory signals. The respiratory phases estimated from different sensors are more consistent, except a global phase shift. Also, the derivatives, which is the instantaneous frequency by our definition, are consistent. The results by CWT with a time-varying spectral band (CWT-tvBPF) and BKD are similar and not shown.

## 4. Material

We consider two real-world databases to demonstrate how to apply the proposed fundamental phase and analysis tools. Particularly, we consider the standardization problem of the notion of phase of a given system when we have multiple time series recorded from one system by different sensors.

### 5.1 First real database—COPD

The first database comes from a prospective study that recruits patients with a clinical diagnosis chronic pulmonary obstructive disease (COPD) according to the Global Initiative for Obstructive Lung Disease Criteria at Chang Gung Memorial Hospital (CGMH), Linkou, Taiwan, from January 2019 to December 2019. The ethics committee of CGMH approved the study. All participants signed informed consent before enrollment. Patients with heart failure (Ejection fraction <40%), lung cancer, atrial fibrillation, using anti-arrhythmic agents and using oxygen were excluded. Total of 63 patients were included, with impaired pulmonary function tests with FVC (Forced vital capacity) 79% (42%-130%) and FEV_1_ (Forced expiratory volume in the 1^st^ second) 58% (21-105%). Experiments were carried out in a quiet room with the temperature maintained at 22-24 °C. Patients were instructed to avoid bronchodilators such as beta-2 agonists, xanthene derivatives, alcohol, and coffee before the experimental test. Before the examination, blood pressure, heart rate, and oxygen saturation were recorded. They wore the pulse oximeter on the finger and ECG electrodes on the chest wall. We recorded the flow signal, impedance, and ETCO_2_ for 5 minutes continuously. To avoid any signal quality issue, the skin was abraded with gel then cleaned with alcohol to reduce skin electrode impedance before ECG electrodes attached, and before the recording, the patient practiced breathing via the flow tube for 45 seconds. Three Actiwave devices were used for recording flow signals and ECG signals. Recording signals were transferred via European Data Format to LabChart 8 analysis software (ADInstruments, USA), and then exported to text files for analysis. The goal is to compare different methods to estimate the phase of the respiratory component, and hence the cardiopulmonary coupling (CPC), from various respiratory sensors.

### 5.2 Second database--LBNP

The second database comes from a previous prospective study that recruits normal subjects for the lower body negative pressure (LBNP) experiment that simulates blood loss. After approval of the protocol by the Yale University Human Committee, healthy subjects of both sexes with age 18-40 years were recruited and provided written consent. Exclusion criteria were current pregnancy or any history of pulmonary, cardiac, or vascular disease including hypertension effectively treated with antihypertensive medication. Subjects were instructed to refrain from alcoholic and caffeinated beverages, strenuous exercise, and tobacco consumption for at least 12 hours prior to the study. A negative pregnancy test was confirmed for all female subjects immediately before participation. Intravenous access was established as a precautionary measure, but fluids were not infused during the study. Thirty-five cases remain in the final analysis. Each subject was monitored with various channels, but we only focused on ECG, finger PPG and PVP (Nonin Medical, Inc., Plymouth, MN). Each procedure was performed in a climate-controlled laboratory at 72°F and 20% humidity. Details of the LBNP experiment are well described [59]. Briefly, after the leg raising test and following a subsequent 5-minute equilibration period, the chamber was sealed to start the LBNP. This entailed 3 minutes phases during which the pressure was progressively decreased by 15 mmHg until either 3 minutes at -75 mmHg were completed or at any phase symptoms consistent with significant hypovolemia occurred. Once an endpoint was reached, pressure in the chamber was increased to -30 mmHg for 1 minute then to 0 for 3 minutes before concluding the study. The goal is to compare different methods to estimate the phase of the hemodynamic component, and hence the instantaneous heart rate (IHR), from PPG and PVP when the hemodynamic is perturbed during the LBNP experiment.

### 5.3 Statistics

For the first real database, the agreement of estimated phases is evaluated by two different approaches. The first one is evaluating the agreement of respiratory phases determined from different sensors. The phase difference is quantified, and for each subject the median and median absolute error are recorded. Results of all subjects are presented as mean±standard deviation. The Mann-Whitney Wilcoxon test is applied to evaluate differences of two variables. P-value less than 0.05 is considered statistically significant. The second one is showing how the phase estimation algorithm influences the widely used algorithm called synchrogram proposed in [60] to quantify CRC [9, 61]. It has been shown in [62] that CRC encodes rich clinical information about COPD patients. The synchrogram is composed of two steps. First, for each subject, determine IHR by the standard cubic spline interpolation of R peak to R peak interval (RRI) time series, where the impact of ventricular premature contraction is modified by the standard RRI correction algorithm [63]. Then, for each subject, quantify the phase lock of the IHR and respiration by the stroboscope idea, which leads to the synchrogram index. The BA plot is applied to evaluate the limit of agreement (LoA) of the synchrogram indices evaluated from different respiratory sensors over the 63 subjects. The bias, LoA, and the associated 95% confidence interval (CI) are reported for each pair of sensors.

For the second real database, we evaluate how the phase of the hemodynamic component, and the associated IHR, could be accurately estimated and the result is independent of the sensor by the proposed fundamental phase and tool. The ground truth IHR is determined by the same method used in the first database, which is sampled at 4Hz. For each subject, the median of the difference of IHR determined by PPG or PVP sampled at 4Hz and the ground truth IHR is recorded and the mean and standard deviation of the median difference by different sensors of all subjects are reported. The Wilcoxon rank test is applied to evaluate if the accuracy of evaluating IHR by different methods is similar, where p<0.05 is considered statistically significant. The BA plot is applied to evaluate the consistency of different sensors, and the same summary statistics as those in the first database are reported.

## 5. Results

### 6.1 First database—COPD

See Figure 2(b) for an illustration of different respiratory signals recorded simultaneously. Physiologically, we know that the inspiration period is shorter than the expiration period, so a respiratory cycle cannot follow a sine wave. Moreover, different respiratory signals capture the pulmonary function from different aspects, so it is natural that different respiratory signals have different WSFs. These facts can be visualized in Figure 2(b), where different respiratory signals oscillate with different WSFs, and all these WSFs are different from a sine wave. Such non-sinusoidal behavior is reflected in the multiples in the spectral domain (see Figure 8(b) for an example). On the other hand, while the lengths and strengths of respiratory cycles vary from cycle to cycle caused by the breathing pattern variability [39], they are similar among different signals since they capture the same physiological system. Therefore, we could model these respiratory signals with the ANHM with *L=1* and different WSFs. We will show that if this polymorphic WSF behavior of respiratory signals is not properly handled, like handled by the traditional method, the phase information might be misleading.

In this database, besides the proposed approaches, we also apply the other commonly applied approaches, including the traditional approaches, like Hilbert transform following a BPF (BPF-HT) and the CWT with a fixed spectral band (CWT-fixed), and ad hoc approaches, like EMD, EEMD and VMD, to estimate the respiratory phase from respiratory signals recorded from different respiratory sensors. We apply the implemented EMD and VMD in Matlab and EEMD implemented in [64] for the reproducibility issue. In EMD, EEMD and VMD, we select the mode that has the highest correlation coefficient with the original respiratory signal, and visually confirm that the selected mode oscillates at the same frequency of the original respiratory signal. We view this selected mode as the reconstructed fundamental component of the respiratory signal. Following the common practice, the phase of the reconstructed fundamental component estimated by Hilbert transform is taken as the estimated phase function. In both BPF-HT and CWT-fixed, we consider the spectral band [0.1,0.4] Hz that covers the usual respiratory rate range, and we view the resulting signal as the fundamental component of the respiratory signal. See Figure 6 for an illustration of the whole procedure of all algorithms applied to different respiratory sensors and the phase difference evaluation. To have a better visual comparison, the reconstructed fundamental components by different methods are shown in Figure 7.

The reconstructed fundamental components of different respiratory signals for the fundamental phase estimation are shown in Figures 4(c) and 5. It is clear that the reconstructed fundamental components by the traditional and ad hoc approaches are non-sinusoidal, while those by the proposed approaches are sinusoidal. Moreover, the reconstructed fundamental components from different respiratory signals are not consistent. For example, the reconstructed fundamental components by EMD vary from one respiratory signal to another. The reconstructed fundamental components by EEMD and VMD are more consistent, but their oscillatory patterns are still not consistent.

In Figure 6(c-e), a further comparison of respiratory phase determined from different respiratory signals and different methods is shown. The decomposed components and the associated phases (shown in Figure 6(c-d) with the phase mod π) by the BPF-HT and CWT-fixed visually look different among different sensors, while those estimated by SST, CWT-tvBPF, and BKD, are similar, except a global phase shift among sensors. Physiologically, this global phase shift describes the intrinsic difference among different respiratory sensors. The phase discrepancy between two sensors is shown in Figure 6(d-e), where the discrepancy fluctuates more in Figure 6(d). This indicates that the phase estimated by CWT-fixed is not consistent if a different sensor is used. Similar results hold for BPF-HT, EMD, EEMD and VMD (not shown).

We further examine the impact of different algorithms on the phase estimation from different sensors by zooming in for details in Figure 8(a). We focus on BPF-HT and SST. It shows that by BPF-HT, the phases of all signals “look close” with wiggles but the IFs, the derivative of the phase, of all channels are different and not physiologically sensible (CWT-fixed result is the same and depicted in Figure 6(d) and not shown here). This shows the problem of applying the traditional approach. The TFRs of those signals by SST are shown in Figure 8(b), where we see a clear and dominant curve around 0.5Hz in each TFR, which means that the IFs of all signals’ fundamental components are consistent. The estimated phases and their derivatives are also consistent among all channels, which are shown in Figure 8(c). Clearly, the results by SST are more sensible and consistent. The results by CWT-tvBPF and BKD are similar but not shown here (See Figure 6(b-c) for an example for CWT-tvBPF and BKD).

After the above visual inspection, we provide quantification of the findings. In Table 1, we report the agreement of phase estimation from different pairs of respiratory sensors and methods. Clearly, the standard deviation is statistically smaller if SST and CWT-tvBPF are applied. On the other hand, note that the mean phase discrepancy might not be zero. This is natural since the mean phase discrepancy reflects the feature of different sensors.

**Table 1.** The median and median absolute error (MAD) of phase difference of all subjects among eight algorithms and three respiratory signals. The p value was evaluated by the Mann-Whitney Wilcoxon test when comparing with results obtained by SST. *: statistically significant with the level 0.05 corrected by the Bonferroni correction. Abbreviation: Impe: Impedance; ETCO2: End tidal CO2; BPF-HT: Hilbert transform followed by a bandpass filter; the Blaschke decomposition (BKD), the empirical mode decomposition (EMD), the ensemble empirical mode decomposition (EEMD), the variational mode decomposition (VMD), CWT-fixed: Continuous wavelet transform with a fixed spectral band; CWT-tvBPF: Continuous wavelet transform with a time varying bandpass filter; SST: Synchrosqueezing transform.

|  |  | Median of phase difference | P value | MAD of phase difference | P value |
| --- | --- | --- | --- | --- | --- |
| Flow VS Impe | BPF-HT | 1.37±0.30 | 0.81 | 0.31±0.10 | <0.00001* |
|  | BKD | 1.30±0.27 | 0.58 | 0.38±0.15 | <0.00001* |
|  | EMD | 1.18±0.29 | 0.001* | 1.05±0.23 | <0.00001* |
|  | EEMD | 1.32±0.29 | 0.91 | 0.45±0.19 | <0.00001* |
|  | VMD | 1.31±0.29 | 0.94 | 0.23±0.17 | 0.0008* |
|  | CWT-fixed | 1.31±0.27 | 0.74 | 0.28±0.20 | <0.00001* |
|  | CWT-tvBPF | 1.31±0.27 | 0.94 | 0.12±0.09 | 0.30 |
|  | SST | 1.32±0.27 | - | 0.12±0.12 | - |
| Flow VS ETCO<sub>2</sub> | BPF-HT | -2.12±0.16 | 0.14 | 0.59±0.14 | <0.00001* |
|  | BKD | -2.15±0.16 | 0.53 | 0.41±0.18 | <0.00001* |
|  | EMD | -1.59±0.47 | <0.00001* | 1.46±0.22 | <0.00001* |
|  | EEMD | -2.12±0.18 | 0.17 | 0.66±0.38 | <0.00001* |
|  | VMD | -2.14±0.15 | 0.29 | 0.32±0.24 | <0.00001* |
|  | CWT-fixed | -2.15±0.15 | 0.56 | 0.39±0.40 | <0.00001* |
|  | CWT-tvBPF | -2.17±0.16 | 0.94 | 0.18±0.31 | 0.02 |
|  | SST | -2.17±0.16 | - | 0.19±0.34 | - |
| Impe VS ETCO<sub>2</sub> | BPF-HT | 1.62±1.70 | <0.001* | 1.83±0.72 | <0.00001* |
|  | BKD | 2.25±1.34 | 0.02 | 1.44±0.82 | <0.00001* |
|  | EMD | 1.28±0.96 | <0.00001* | 1.86±0.29 | <0.00001* |
|  | EEMD | 2.05±1.36 | 0.0001 | 1.42±0.73 | <0.00001* |
|  | VMD | 2.03±1.56 | 0.01 | 1.16±0.93 | 0.002* |
|  | CWT-fixed | 2.12±1.42 | 0.003 | 1.22±1.01 | <0.00001* |
|  | CWT-tvBPF | 2.34±1.39 | 0.91 | 0.61±0.77 | 0.03 |
|  | SST | 2.32±1.45 | - | 0.55±0.71 | - |

To further demonstrate its clinical application, the CRC indices of 63 subjects determined by the synchrogram from different respiratory signals are compared by the Bland-Altman plot. The bias, LoA, and the associated 95% CI are reported for each pair of sensors in Table 2. The results suggest that if the respiratory phase is estimated by SST, CWT-tvBPF or BKD, the CRC index is less impacted by the sensor type compared with other methods.

**Table 2:** Summary of Bland-Altman (BA) plot of the cardiopulmonary coupling index determined from different phase reconstruction methods, including the Hilbert transform followed by a bandpass filter (BPF-HT), the Blaschke decomposition (BKD), the empirical mode decomposition (EMD), the ensemble empirical mode decomposition (EEMD), the variational mode decomposition (VMD), the continuous wavelet transform (CWT) with a fixed spectral band (CWT-fixed), the CWT with the proposed time-varying bandpass filter (CWT-tvBPF) and the synchrosqueezing transform (SST), and different respiratory signals, including flow, impedance (Impe) and End tidal CO_2_ (ETCO_2_). The comparison suggests that the agreements of respiratory phases determined by CWT-tvBPF and SST are higher than other methods. CI means confidence interval.

| Algorithm | Signals under Comparison | Bias | Upper 95% agreement bound [95% CI] | Lower 95% agreement bound [95% CI] |
| --- | --- | --- | --- | --- |
| BPF-HT | Flow VS Impe | 0.01 | 0.11 [0.09, 0.13] | -0.08 [-0.06, -0.10] |
|  | Flow VS ETCO<sub>2</sub> | 0.04 | 0.23 [0.19, 0.28] | -0.16 [-0.11, -0.20] |
|  | Impe VS ETCO<sub>2</sub> | 0.02 | 0.17 [0.14, 0.21] | -0.13 [-0.09, -0.16] |
| EMD | Flow VS Impe | -0.07 | 0.25 [0.18, 0.33] | -0.40 [-0.33, -0.48] |
|  | Flow VS ETCO<sub>2</sub> | -0.14 | 0.26 [0.17, 0.35] | -0.54 [-0.46, -0.64] |
|  | Impe VS ETCO<sub>2</sub> | -0.07 | 0.25 [0.18, 0.32] | -0.39 [-0.32, -0.46] |
| EEMD | Flow VS Impe | -0.08 | 0.24 [0.17, 0.31] | -0.41 [-0.34, -0.48] |
|  | Flow VS ETCO<sub>2</sub> | -0.007 | 0.14 [0.11, 0.18] | -0.15 [-0.12, -0.19] |
|  | Impe VS ETCO<sub>2</sub> | 0.08 | 0.39 [0.32, 0.46] | -0.23 [-0.17, -0.31] |
| VMD | Flow VS Impe | 0.007 | 0.11 [0.14, 0.09] | -0.09 [-0.08, -0.12] |
|  | Flow VS ETCO<sub>2</sub> | 0.02 | 0.13 [0.16, 0.11] | -0.09 [-0.07, -0.12] |
|  | Impe VS ETCO<sub>2</sub> | 0.01 | 0.08 [0.09, 0.06] | -0.05 [-0.04, -0.07] |
| BKD | Flow VS Impe | 0.0003 | 0.04 [0.05, 0.006] | -0.04 [-0.005, -0.05] |
|  | Flow VS ETCO<sub>2</sub> | -0.003 | 0.03 [0.04, 0.02] | -0.03 [-0.007, -0.04] |
|  | Impe VS ETCO<sub>2</sub> | -0.02 | 0.08 [0.10, 0.06] | -0.12 [-0.09, -0.14] |
| CWT-fixed | Flow VS Impe | 0.13 | 0.11 [0.09, 0.13] | -0.08 [-0.06, -0.10] |
|  | Flow VS ETCO<sub>2</sub> | 0.01 | 0.11 [0.09, 0.13] | -0.09 [-0.07, -0.11] |
|  | Impe VS ETCO<sub>2</sub> | -0.004 | 0.08 [0.06, 0.10] | -0.09 [-0.07, -0.10] |
| CWT-tvBPF | Flow VS Impe | 0.0003 | 0.04 [0.04, 0.05] | -0.04 [-0.03, -0.05] |
|  | Flow VS ETCO<sub>2</sub> | -0.003 | 0.03 [0.02, 0.04] | -0.04 [-0.03, -0.05] |
|  | Impe VS ETCO<sub>2</sub> | -0.003 | 0.04 [0.03, 0.04] | -0.04 [-0.03, -0.05] |
| SST | Flow VS Impe | 0.0005 | 0.04 [0.03, 0.05] | -0.05 [-0.04, -0.06] |
|  | Flow VS ETCO<sub>2</sub> | -0.004 | 0.04 [0.04, 0.06] | -0.05 [-0.04, -0.06] |
|  | Impe VS ETCO<sub>2</sub> | 0.002 | 0.04 [0.04, 0.06] | -0.04 [-0.03, -0.06] |

### 6.2 Second database--LBNP

Based on the findings in the first database, we only consider SST and BPF-HT in this database to simplify the discussion. In the past decades, researchers have established a lot of physiological knowledge about PPG [45] and PVP [65]. PPG reflects the volume change during the cardiac cycle, and PVP is the pressure recorded from the peripheral venous. It is well known that the cardiac dynamics during the systolic and diastolic phases are different, which induce the non-sinusoidal oscillatory patterns of PVP and PPG. On the other hand, since PPG and PVP both reflect the cardiac cycles, they share the same instantaneous frequency. Thus, modelling PVP and PPG with different WSFs in the ANHM is justified.

Two segments of PPG and PVP simultaneously recorded from a healthy subject during the baseline and during the -75mmHg pressure (simulating a 1,500cc blood loss) are shown in Figure 2(c). For a single subject, the WSFs of PPG and PVP are different in different physiological statuses. Based on the COPD result, we only compare BPF-HT and SST. In Figure 9(a), we show the estimated phases by BPF-HT with different spectral bands, and both are not sensible.

**Figure 9:**
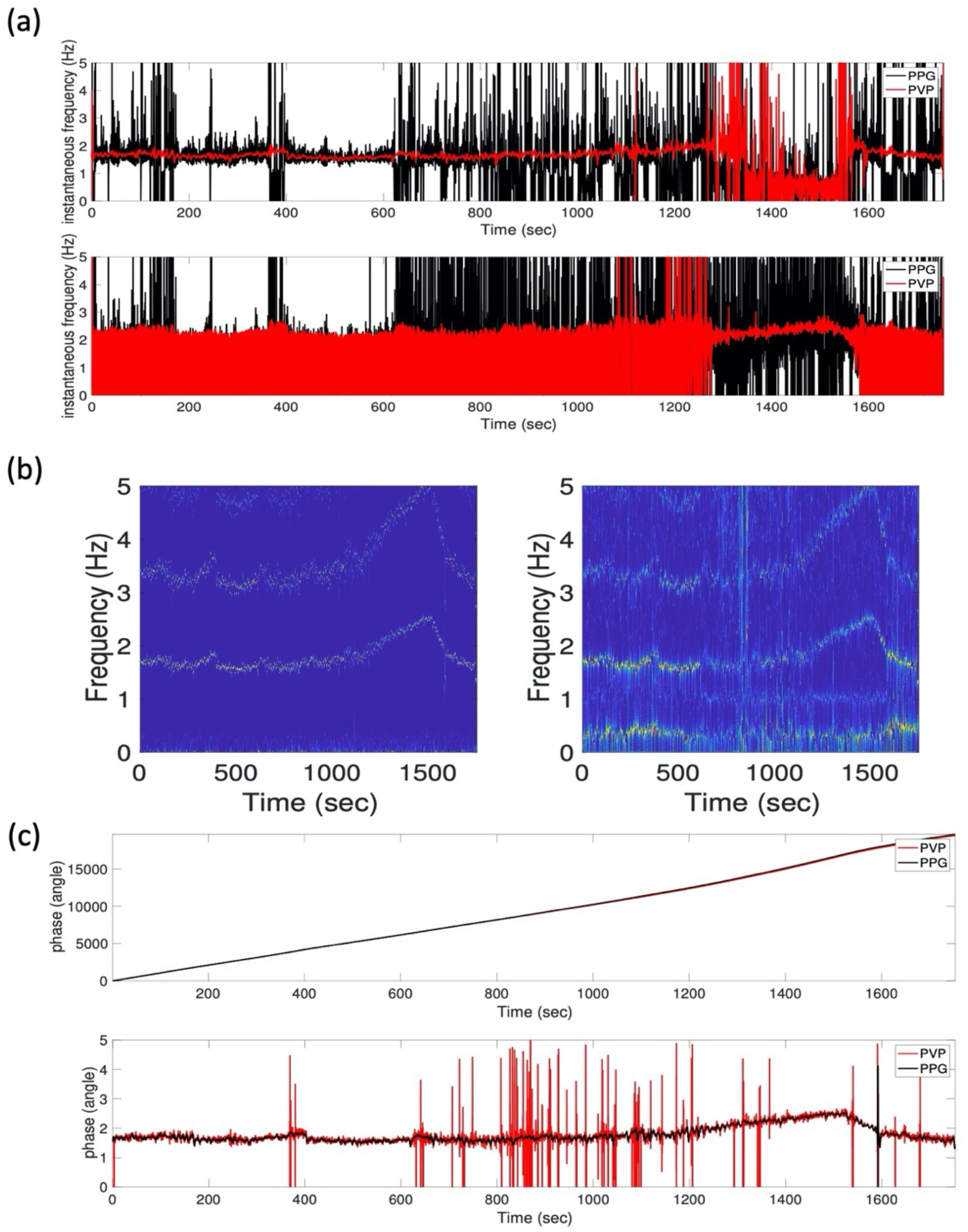
(a) An example of estimating phases of photoplethymogram (PPG) and peripheral venous pressure (PVP) signals by the Hilbert transform followed by a band-pass filter (BPF-HT) with different spectral bands, where the derivatives of the estimated phases, the instantaneous frequencies, are shown for a comparison. In the top panel, the spectral band is 0.5-2 Hz, while in the bottom panel, the spectral band is 0.5-4 Hz. The lower spectral bound 0.5 Hz is chosen to avoid the respiratory component impact, and the upper spectral bound 2 or 4 Hz is chosen to include sufficient cardiac information. However, in both cases, the phases, and hence their derivatives, are not consistent between the PPG and PVP signals. The difference is clear over 1300-1600 seconds. This comes from the fact that the heart rate increases dramatically during 1300-1600 seconds during the lower body negative pressure experiment, and the chosen spectral bands, if fixed, may include insufficient fundamental component information or 14 may be contaminated by its multiples. (b) An example of the time-frequency 15 representations (TFRs) determined by the synchrosqueezing transform (SST). The TFR of the photoplethymogram (PPG) is shown on the left and the TFR of the peripheral venous pressure (PVP) is shown on the right. (c) the phases and the associated instantaneous frequency of the cardiac component in the PPG and PVP signals. The phases associated with the PPG and PVP signals are shown on the top, and the associated instantaneous frequency of the hemodynamic oscillation determined by differentiating the estimated phases are shown in the bottom. The phases of PPG and PVP signals are consistent. Also, the derivatives, which is the instantaneous frequency by our definition, are consistent, except for some unwanted spikes in the PVP signal. This comes from the well-known fact that the PVP signal is usually too noisy to analyse with traditional tools

See Figure 9(b) for the TFRs of PPG and PVP by SST. In both TFRs, there is a dominant curve starting at around 1.8Hz, surging to 2.5Hz in the 4/5 of the experiment (around -75mmHg), and then coming back to 2Hz. This reflects the fact that during a massive blood loss, tachycardia happens. The multiple can also be clearly seen. There are more structures in the TFR of PVP, like a dominant curve at 0.3Hz, and a vague curve around 1Hz from 600sec to 1100sec. The curve at 0.3Hz is associated with the respiration, and the curve around 1Hz from 600sec to 1100sec is unknown to our knowledge and needs more exploration. These indicate that there are multiple non-harmonic IMT components constituting PVP. In Figure 9(c), the proposed tvBPF successfully gives a sensible phase of the cardiac component, and hence the IHR via differentiation. Since both PPG and PVP reflect the cardiac cycles, it makes sense that the estimated IHRs are close. However, the IHR estimated from PVP is noisy. It is not surprising since PVP is usually noisy.

Quantitatively, among all subjects, the median IHR estimation errors from PPG and PVP by SST are 0.0007±0.0004 Hz and 0.0077±0.0049 Hz respectively, while the median IHR estimation errors from PPG and PVE by BPF-HT are 1.18±0.185 Hz and 1.165±0.176 Hz respectively. The p-value of the Wilcoxon test comparing the performance of BPF-HT and SST of estimating the IHR from PPG and PVP are <10^−4^ and <10^−4^ respectively, which are both statistically significant after the Bonferroni correction. The agreement bias of estimating the IHR from PPG and PVP by SST and BPF-HT are 0.006 and -0.019 respectively, with the upper LoA 1.34 [95% CI: [1.33, 1.34]] and 0.042 [95% CI: [0.041, 0.042]], and the lower LoA -1.32 [95% CI: [-1.32, - 1.33]] and -0.081 [95% CI: [-0.080, -0.081]] respectively. The above results indicate that SST could estimate IHR more accurately compared with BPF-HT from both PPG and PVP.

## 6. Discussion

Phase is a fundamental quantity describing an oscillatory signal. We show that the phase estimated by the traditional approaches based on the analytic function model and ad hoc approaches without theoretical support might not be suitable for biomedical oscillatory signals. We then suggest standardizing the phase by modeling oscillatory biomedical time series by ANHM and consider the proposed fundamental phase. Then, we suggest applying the tvBPF scheme with time-frequency analysis tools, like SST and CWT, or BKD, to extract the fundamental phase. We demonstrate one important application of the proposed scheme to standardize the notion of phase from different signals recorded from the same system with different sensors in two real-world databases. We show that the proposed fundamental phase, when extracted with a proper algorithm, is less independent of the sensor type. We also provide a Matlab implementation for researchers.

This result is critical in the heterogeneous clinical environment, which becomes more diverse in the post-COVID 19 era. Take respiratory signal as an example. In digital medicine, different respiratory sensors are available to acquire respiratory information, while they differ from different perspectives, like signal quality, ease-of-use and what information could be acquired. For example, a continuous recording of flow signals is less suitable for home-based healthcare, while it provides more information. On the other hand, the impedance respiratory signal could be easily acquired, but the quality is less stable and less information, like tidal volume, could be directly obtained. Without a standardized approach, it would be challenging to exchange information if different respiratory sensors are used, and an advance of scientific knowledge could be limited. We would expect that if this fundamental phase is properly used, it helps scientific research from different aspects, including information sharing, discussion of the scientific findings among different equipment, and others.

We shall mention technical details of algorithms other than SST and CWT. While EMD and its variations like EEMD and VMD have been widely applied, to our best knowledge, there does not exist sufficient theoretical support for how they achieve a proper decomposition of signals satisfying ANHM so that the phase can be properly estimated and in practice we have found several potential problems that we cannot fix without having more insights into the algorithm. On the other hand, as a novel TF analysis tool based on the Blaschke decomposition in complex analysis, we have several theoretical supports about how it works and we know when it is limited. For example, the cardiopulmonary coupling index shown in Table 2 determined by BKD is as consistent among different sensors as those obtained by SST and CWT-tvBPF. This comes from the fact that the fundamental components of all respiratory signals considered in this work dominate their harmonics. However, when this fact does not hold, a modification of the BKD algorithm and a new theoretical support is needed. In general, when there are multiple oscillatory components in the signal, how to apply BKD is still open. We shall emphasize that the poor numerical results of BKD in this paper are expected from the theoretical perspective, and it does not indicate that BKD is not useful for other applications.

To achieve a fully automatic estimation of the fundamental phase, there are several challenges we shall resolve. First, Definition 1 does not capture more complicated dynamics; for example, the wax-and-wane pattern that an oscillatory component might only appear in a subinterval but not the whole recorded signal. For example, the respiratory component in PPG might exist for a while and disappear over another period. How to automatically determine when a non-harmonic IMT function exists could be framed as a change point problem. While some work has been developed recently, more work is needed. Note that to avoid complications and focus on the notion of fundamental phase, we focus on (5) in this paper. On the other hand, it is scientifically critical to ask if a collected signal can be well modeled by the ANHM. For the signals we analyzed as examples in this paper, the respiratory signals, PPG and PVP, we have clear physiological knowledge to support why the ANHM is proper. However, when we lack background knowledge, or when the signal is noisy so that we cannot identify the oscillatory pattern, a quantitative assessment of the chosen model is critical. To our knowledge, this problem is so far still open, particularly for signals with time-varying frequency and amplitude with non-sinusoidal oscillation.

Second, notice that the discussion in this paper is specifically for the oscillatory signals with time-varying frequency and amplitude with non-sinusoidal oscillatory pattern. In general, it might not be directly applied to model stochastic-process-like physiological signals, like R peak to R peak interval (RRI) time series for heart rate variability analysis, EEG or local field potential. Take EEG as an example. It is a common practice to define spectral bands based on prior knowledge; for example, the high frequency and low frequency bands for the RRI time series analysis and alpha, beta, delta, gamma and theta bands commonly considered in the EEG analysis. Then, find the phase of the spectral band of interest and discuss how it changes or is coupled with other bands according to time. Since the signal associated with a given spectral band is a broadband signal, modeling its phase by Definition 1 considered for oscillatory signals is not proper. Since a random process has a fundamentally different behavior compared with oscillatory signals we consider in this paper, a systematic discussion of the phase of these stochastic-process-like signals shall be further explored.

Third, the ridge extraction algorithm (12) might not be the optimal solution to extract IF, particularly when the multiples have stronger strength, or even when the fundamental component does not exist. The solution is combining rectification and the de-shape algorithm [48]. Since the focus of this paper is phase, these topics are not extensively discussed and we refer readers with interest in this solution to [48].

Fourth, in practice we may or may not know *L* in (5). Usually, we have background knowledge guiding us about *L*. See the PPG and respiratory signal as biomedical examples shown in Section 2. However, when we do not have sufficient background knowledge, a rigorous estimate of *L* from the function *f*(*t*) is important for the scientific purpose. There are some ad hoc approaches toward this goal; for example, via the de-shape algorithm [48] and the iterative reconstruction process. In short, the de-shape algorithm could help eliminate the impact of non-sinusoidal oscillation in the time-frequency domain, so that we can count the number of components at each time. With the estimated *L*, then, reconstruct the associated oscillatory components. Finally, subtract the estimated oscillatory components from the signal and estimate the stationarity or the properties of the preassigned noise model to establish how confident we are about the estimated *L*. We should mention that while we could apply this approach to determine *L*, a systematic approach with confidence and theoretical support is so far a challenging problem, and to our knowledge, it is still an open challenging problem in the field. Since estimating *L* is itself an independent work and is out of the scope of this paper, we leave it to our future work. Tackling the above challenges will be our future work.

## 7. Conclusions

The proposed combination of ANHM and tvBPF with TF analysis tools gives a meaningful phase information and alleviates the challenges of nonuniform phase extraction across different recording modalities. Under the existing theoretical supports, we expect its scientific impact on a broad range of applications.

## Acknowledgements

The authors would like the thank the anonymous reviewers for their constructive and valuable comments and critiques that highly increases the quality of this paper. HT Wu would like to thank Dr. Yu-Chen Huang for her help in preparing Figures 6 and 7.

## Availability of data and materials

The code used in this paper are available in https://github.com/hautiengwu/ReconsiderPhase

## References

1. H Gesche, D Grosskurth, G Küchler, A Patzak, Continuous blood pressure measurement by using the pulse transit time: comparison to a cuff-based method, Eur. J. Appl. Physiol., 112 (2012), 309--315.

2. SH Kim, JG Song, JH Park, JW Kim, YS Park, GS Hwang, Beat-to-beat tracking of systolic blood pressure using noninvasive pulse transit time during anesthesia induction in hypertensive patients, Anesth. Analg. 116 (2013), 94--100

3. C Schäfer, MG Rosenblum, HH Abel, J Kurths, Synchronization in the human cardiorespiratory system, Phys. Rev. E. 60 (1999), 857.

4. M Weiss, G Weiss, A derivation of the main results of the theory of hp spaces, Rev. de la Union Mat. Argentina, 20 (1962), 63--71.

5. RP Bartsch, AY Schumann, JW Kantelhardt, T Penzel, PC Ivanov, Phase transitions in physiologic coupling, Proc. Natl. Acad. Sci. U.S.A., 109(2012), 10181--10186.

6. D Dvorak, AA Fenton, Toward a proper estimation of phase–amplitude coupling in neural oscillations, J. Neurosci. Methods, 225 (2014), 42--56.

7. D Khodagholy, JN Gelinas, G Buzsáki, Learning-enhanced coupling between ripple oscillations in association cortices and hippocampus, Science, 358 (2017), 369--372.

8. JP Lachaux, E Rodriguez, J Martinerie, C Adam, D Hasboun, FJ Varela, A quantitative study of gamma-band activity in human intracranial recordings triggered by visual stimuli. European Journal of Neuroscience, 12 (2000), 2608--2622.

9. MG Rosenblum, L Cimponeriu, A Bezerianos, A Patzak, R Mrowka, Identification of coupling direction: application to cardiorespiratory interaction, Phys. Rev. E., 65 (2002), 041909.

10. MG Rosenblum, AS Pikovsky, Detecting direction of coupling in interacting oscillators, Phys. Rev. E., 64 (2001), 045202.

11. MR Miller, J Hankinson, V Brusasco, F Burgos, R Casaburi, A Coates, R Crapo, P Enright, C Van Der Grinten, P Gustafsson, Standardisation of spirometry, Eur. Respir. J., 26 (2005), 319--338.

12. R Klabunde, “Cardiovascular physiology concepts,” L.W.W., 2011.

13. AL Hodgkin, AF Huxley, A quantitative description of membrane current and its application to conduction and excitation in nerve, J. Physiol.,117 (1952), 500--544.

14. T Wigren, Model order and identifiability of non-linear biological systems in stable oscillation, IEEE/ACM, Trans, Comput, Biol, Bioinform., 12 (2015), 1479--1484.

15. P Ashwin, S Coombes, R Nicks, Mathematical frameworks for oscillatory network dynamics in neuroscience, J. Math. Neurosci., 6 (2016), 1--92.

16. J Keener, J Sneyd, “Mathematical physiology 1 Cellular physiology”, Springer, 2009.

17. JD Murray, “ Mathematical biology: I. An introduction, “ Springer Science & Business Media, 2007.

18. B Picinbono, On instantaneous amplitude and phase of signals, IEEE Trans. Signal. Process., 45 (1997), 552--560.

19. B Van der Pol B, The fundamental principles of frequency modulation, J. Inst. Electr. Eng. 93 (1946), 153--158.

20. A Rihaczek, E Bedrosian. Hilbert transforms and the complex representation of real signals. Proc. IEEE., 54 (1966) 434--435.

21. D Gabor, Theory of communication. Part 1: The analysis of information. J. IEEE., 93 (1946), 429--441.

22. E Bedrosian. A product theorem for Hilbert transforms, Proc. IEEE., 5 (1962), 868--869

23. R Nevanlinna, “The First Main Theorem in the Theory of Meromorphic Functions. Analytic Functions.” Springer, 1970.

24. AV Oppenheim, “ Discrete-time signal processing,” Pearson Education India, 1999.

25. M Feldman M, Time-varying vibration decomposition and analysis based on the Hilbert transform, J. Sound. Vib., 295 (2006), 518--530.

26. I Daubechies, Ten lectures on wavelets. SIAM, Philadelphia, (1992). MR 93e 42045.

27. L Cohen. Time-frequency distributions-a review, Proc. IEEE., 77 (1989) 941--981.

28. P Flandrin, Time-frequency/time-scale analysis. Academic press (1998)

29. MR Nahon, “Phase evaluation and segmentation,” Spirnger, 2001.

30. T Qian, Intrinsic mono-component decomposition of functions: an advance of Fourier theory, Math. Methods. Appl. Sci., 33 (2010), 880–891.

31. NE Huang, Z Shen, SR Long, MC Wu, HH Shih, Q Zheng, NC Yen, CC Tung, HH Liu, The empirical mode decomposition and the Hilbert spectrum for nonlinear and non-stationary time series analysis, Proc. R. Soc. Lond., 454 (1998), 903--995.

32. Z Wu, NE Huang, Ensemble empirical mode decomposition: a noise-assisted data analysis method. Advances in adaptive data analysis, 1 (2009) 1--41.

33. K Dragomiretskiy, Z Dominique, Variational mode decomposition, IEEE Trans. signal processing, 62 (2013) 531--544.

34. M Chavez, M Besserve, C Adam, J Martinerie, Towards a proper estimation of phase synchronization from time series, J. Neurosci. Methods., 154 (2006), 149--160.

35. M Le Van Quyen, J Foucher, JP Lachaux, E Rodriguez, A Lutz, J Martinerie, FJ Varela, Comparison of Hilbert transform and wavelet methods for the analysis of neuronal synchrony. J. Neurosci. Methods 2001(111) 83--98.

36. HT Wu, Current state of nonlinear-type time-frequency analysis and applications to high-frequency biomedical signals, Curr. Opin. Syst. Biol., 23(2020), 18--21

37. YC Chen, MY Cheng, HT Wu, Non-parametric and adaptive modelling of dynamic periodicity and trend with heteroscedastic and dependent errors, J. R. Stat. Soc. Series. B Stat. Methodol., (2014), 651--682.

38. I Daubechies, J Lu, HT Wu, Synchrosqueezed wavelet transforms: An empirical mode decomposition-like tool, Appl. Comput. Harmo. Anal. 30 (2011), 243--261

39. G Benchetrit, Breathing pattern in humans: diversity and individuality, Respir. Physiol., 122 (2000), 123--129.

40. YT Lin, HT Wu, J Tsao, HW Yien, SS Hseu, Time-varying spectral analysis revealing differential effects of sevoflurane anaesthesia: non-rhythmic-to-rhythmic ratio, Acta. Anaesthesiol., 58(2014), 157--167.

41. D Vakman, On the analytic signal, the Teager-Kaiser energy algorithm, and other methods for defining amplitude and frequency, IEEE Trans. Signal. Process. 44 (1996), 791--797.

42. J Huang, Y Wang, L Yang, Vakman’s problem and the extension of Hilbert transform. Applied and computational harmonic analysis, 34 (2013), 308--316.

43. D Vakman, On the definition of concepts of amplitude, phase and instantaneous frequency of a signal, Radio. Eng. Electron. Phys. 17 (1972), 754--759.

44. A Nuttall, E Bedrosian, On the quadrature approximation to the Hilbert transform of modulated signals, Proc. IEEE., 54 (1966), 1458–1459.

45. KH Shelley, Photoplethysmography: beyond the calculation of arterial oxygen saturation and heart rate. Anesth. Anal., 105 (2007), 31--36.

46. S Luo, WJ Tompkins, JG Webster, Cardiogenic artifact cancellation in apnea monitoring. Proceedings of 16th Annual International Conference of the IEEE Engineering in Medicine and Biology Society, IEEE. 2(1994), 968--969.

47. HT Wu. 2013, Instantaneous frequency and wave shape functions (I), Appl. Comput. Harmon. Anal., 35 (2013), 181--199.

48. CY Lin, L Su, HT Wu, Wave-shape function analysis, J. Fourier Anal. Appl., 24 (2018), 451--505.

49. SM Sourisseau, HT Wu, Z Zhou, Inference of synchrosqueezing transform--toward a unified statistical analysis of nonlinear-type time-frequency analysis, arXiv preprint 1904.09534, 2019.

50. YT Lin, J Malik. HT Wu, Wave-shape oscillatory model for nonstationary periodic time series analysis, Foundation of Data Science, 3(2021), 99–131.

51. Code for this paper. https://github.com/hautiengwu/ReconsiderPhase

52. HT Wu, Adaptive analysis of complex data sets. Princeton University, 2011

53. WAVELAB 850, https://statweb.stanford.edu/~wavelab/.

54. The Time-frequency tool box. http://tftb.nongnu.org.

55. Garnett, J., Bounded analytic functions. Vol. 236. 2007: Springer Science & Business Media.

56. RR Coifman, S Steinerberger, HT Wu, Carrier frequencies, holomorphy, and unwinding, SIAM J. Math. Anal., 49 (2017), 4838--4864.

57. Coifman, R.R. and S. Steinerberger, Nonlinear phase unwinding of functions. Journal of Fourier Analysis and Applications, 2017. 23(4): p. 778–809.

58. BKD code. https://github.com/hautiengwu/BlaschkeDecomposition

59. AA Alian, NJ Galante, NS Stachenfeld, DG Silverman, KH Shelley, Impact of central hypovolemia on photoplethysmographic waveform parameters in healthy volunteers part 2: frequency domain analysis, J. Clin. Monit. Comput., 6 (2011), 387--396.

60. C Schäfer, MG Rosenblum, J Kurths, HH Abel, Heartbeat synchronized with ventilation, nature, 392 (1998), 239--240.

61. AS Pikovsky, MG Rosenblum, GV Osipov, J Kurths, Phase synchronization of chaotic oscillators by external driving, Physica. D., 104 (1997), 219--238.

62. YC Huang, TY Lin, HT Wu, PJ Chang, CY Lo, TY Wang, CH Kuo, SM Lin, FT Chung, HC Lin, YL Lo, Cardiorespiratory coupling is associated with exercise capacity in patients with chronic obstructive pulmonary disease, BMC. Pulm. Med., 21 (2021)

63. M Peltola. Role of editing of RR intervals in the analysis of heart rate variability, Front. Physiol., 3(2012), 148.

64. EMD codes : https://github.com/benpolletta/HHT-Tutorial/tree/master/HuangEMD.

65. R Wardhan, K Shelley, Peripheral venous pressure waveform. Curr Opin Anaesthesiol. 22 (2009), 814–21.

66. I Daubechies, Y Wang, HT Wu, ConceFT: Concentration of frequency and time via a multitapered synchrosqueezed transform. Philosophical Transactions of the Royal Society A: Mathematical, Physical and Engineering Sciences 374.2065 (2016): 20150193.

67. S Meignen, DH Pham, S McLaughlin, On demodulation, ridge detection, and synchrosqueezing for multicomponent signals, IEEE Trans. Signal Process., 65:8 (2017), 2093–2103.

68. MA Colominas, HT Wu, Decomposing non-stationary signals with time-varying wave-shape functions. IEEE Trans. Signal Process. 69 (2021), 5094–5104.

